# Microfluidic cultivation enables probiotic growth under thermal stress

**DOI:** 10.64898/2026.10.23.761394

**Authors:** Ali Karimi, Mohammad Yaghoobi, Shiying Zhang, Genevieve Flock, Gianna P. Ahnrud, Patrick Marek, Kenneth W. Simpson, Alireza Abbaspourrad

**Affiliations:** Department of Food Science, College of Agricultural and Life Sciences, Cornell University, Stocking Hall, Ithaca, NY, 14853 USA; Department of Clinical Sciences, College of Veterinary Medicine, Cornell University, Ithaca, NY, 14853 USA; DEVCOM Soldier Center, Soldier Sustainment Directorate, Combat Feeding Division, Food Protection & Innovative Packaging Team, Natick, MA, 01760 USA

**Keywords:** Microfluidics, Continuous bacterial cultivation, Thermal adaptation, Stress response, Probiotics

## Abstract

Thermal stress tolerance is essential for preserving the viability and functionality of probiotic bacteria. Conventional batch cultures offer limited control over temperature changes across space and time. We have introduced an integrated microfluidic platform to continuously cultivate bacteria within a precisely defined and tunable thermal environment, and used *Lacticaseibacillus rhamnosus* as a model system. The device features serially connected microchambers with independently controlled heating zones, creating distinct and dynamic thermal profiles as the bacteria travel from one microchamber to the next. Biomass accumulation was observed at 37 °C within 48 h in all microchambers, no growth occurred at 49 °C even after 72 h. When the cells were exposed to a periodic thermal landscape, where an increase in temperature in one chamber was followed by a return to 37 °C in the next, growth was observed in the last chamber (49 °C) after three days indicating adaptation to thermal stress. Cells that grew on-chip at 49 °C showed no growth in off-chip experiments. Continuous monitoring of pH at the chip outlet confirmed ongoing metabolic activity in growing populations. Tailored microfluidic thermal environments were shown to help bacteria grow even under otherwise inhibitory conditions.

## 1 Introduction

Probiotic bacteria are live microorganisms that, when consumed in adequate amounts, confer health benefits on the host, including support for the gut, immune, and metabolic health (Kechagia et al., 2013). Probiotics can lose viability during processes such as drying, pasteurization, and transporting in non-refrigerated conditions, compromising their health-promoting properties (Iaconelli et al., 2015; Wendel, 2022). Therefore, understanding the response of probiotics to thermal stress is essential for expanding their use in food and beverage applications (Sanders et al., 1999; De Angelis et al., 2004; Prasad et al., 2003; Zhen et al., 2020).

Temperature is a crucial factor during the proliferation of microorganisms (Chen and Shakhnovich, 2010); elevated temperatures increase protein denaturation and misfolding (Chen and Shakhnovich, 2010), can cause ribosome and RNA destabilization (Jaquet et al., 2022), and alter membrane fluidity (de Mendoza and Cronan, 1983). Despite protective systems such as heat-shock proteins (HSPs) (Hu et al., 2022), bacterial responses to thermal stress vary substantially depending on the surrounding environmental conditions and the specific strain (Eng Ashley Y. et al., 2023). The limited number of strategies available to protect probiotics from high temperatures highlights the importance of studying thermal adaptation under spatially and thermally well-controlled environments (Nagy et al., 2018).

Assessment of bacterial response to a thermally stressed environment has been done as transient heat-shock and long-term exposure. Transient heat-shock treatments are used to investigate the short-term physiological responses of bacterial cells to sudden increases in temperature (Morita et al., 1999; De Angelis et al., 2004; Zhen et al., 2020). In contrast, long-term experiments, such as adaptive laboratory evolution (ALE), grow bacteria under thermally optimal (Lenski et al., 1991) or thermally stressed (Leroi et al., 1994; Bennett and Lenski, 2007) conditions. ALE experiments rely on prolonged cultivation times to allow beneficial mutations to emerge and remain in the population based on natural selection (Dragosits and Mattanovich, 2013). The duration of ALE experiments ranges from several days (Bennett et al., 1990; Rudolph et al., 2010) to more than a decade (Couce et al., 2024). The outcome of experimental thermal adaptation can arise from genetic mutations that become fixed in the population (Riehle et al., 2003; Deatherage et al., 2017) or from non-genetic phenotypic changes induced by environmental conditions, often regarded as acclimation (Bennett and Lenski, 1997).

Long-term thermal adaptation studies are conventionally performed by serial transfer of batch cultures to elevated temperatures over multiple generations (Dragosits and Mattanovich, 2013). Although this method is straightforward (Lenski, 2023), it is time-consuming, labor intensive, requires large volumes of medium, and runs the risk of cross-contamination. In addition, the periodic removal of flasks from the incubator for sampling introduces unintended temperature fluctuations, which compromise the accuracy of the observed responses. Alternative platforms include chemostats, in which cells are maintained at a constant growth rate through continuous replenishment of fresh nutrients and removal of culture medium (Gresham and Dunham, 2014; Huang et al., 2023). The loss of a robust stress response, however, is a common outcome of chemostat selection (Gresham and Dunham, 2014). Despite offering automation, chemostats are often more costly to set up and suffer from operation complexity and potential biofilm formation on the container walls (Lenski, 2023).

Microfluidic systems offer precise control over spatial gradients, fluid flow, and localized stress conditions (Zoheir et al., 2024), while enabling the automation of adaptation studies (Wong et al., 2018). As an example, microdroplet platforms enabled high-throughput cultivation and screening of bacteria under chemical stressors, such as antibiotics (Jakiela et al., 2013; Postek and Garstecki, 2022; Agnihotri et al., 2025). However, microdroplet platforms are often only suitable for short-term cultivation and cannot be used in high-temperature conditions due to the risk of liquid evaporation. Connected microchambers with individually controlled stressor magnitude have also been used to generate segregated chemical gradients, exposing the cells to stressed environments to study the adaptive evolution of motile (Zoheir et al., 2021) and immotile (Nguyen et al., 2024; Li et al., 2026) bacteria. The movement of cells between chambers using fluid flow resembles serial batch transfer, while the continuous nutrient supply is analogous to a chemostat. However, the generation of a thermally stressed landscape in a continuous-flow microfluidic system has not yet been addressed.

Here, we developed an integrated microfluidic platform to study how bacteria adapt to thermally stressed environments during continuous cultivation. The system consists of a series of connected microchambers with independently controlled heating zones, allowing us to maintain well-defined thermal conditions while continuously supplying nutrients and removing waste. By combining thermoelectric control with real-time imaging, the platform enables continuous monitoring of bacterial growth and metabolic activity. Using this approach, we explored bacterial responses across different thermal landscapes, including uniform, gradient, and periodic conditions, and found that dynamic thermal environments on a microfluidic platform can enable growth under otherwise growth-inhibiting temperatures. Together, this work highlights the potential of microfluidic systems for studying microbial adaptation with precise spatial and temporal control.

## 2. Methods

### 2.1 Bacterial culture and maintenance

*Lacticaseibacillus rhamnosus* (Hansen), purchased from American Type Culture Collection (Gaithersburg, MD, USA), was streaked to a de Man, Rogosa, Sharpe (MRS) agar plate (Sigma-Aldrich, MO, USA) and incubated at 37 °C for 48 h. A single colony was picked from the plate and inoculated into 5 mL of sterilized MRS broth (BD Difco, NJ, USA) and incubated overnight at 37 °C. Monoclonal stock aliquots were prepared in 20 v/v% glycerol and stored at –80 °C. To examine the viability of the cultures, we used the pour plating technique and enumerated the colony-forming units (CFUs).

### 2.2 Off-chip assessment of thermal response

Cultures were prepared in a reduced evaporation 96-well plate (CELLTREAT, Ayer, MA, USA) using cryopreserved inocula at cell densities of 10^6^ or 10^7^ CFU mL^−1^. The optical density at 600 nm of each well was measured using a microplate reader (SpectraMax iD3 Multi-Mode, Molecular Devices, USA) at a predefined temperature every 15 min. We obtained the growth parameters, maximum optical density (A), maximum specific growth rate (μ_max_), and growth lag duration (λ), from fitting the data to the reparametrized Gompertz model (Zwietering et al., 1990) using a MATLAB script. Nonlinear least-squares optimization was used to fit the model to each dataset, and goodness-of-fit was evaluated using the coefficient of determination (R^2^) (Supplementary Information **Note 1**).

### 2.3. Microfluidic setup fabrication

Conventional soft lithography method was used for fabrication of the microfluidic device (Xia and Whitesides, 1998). Briefly, the master mold for the microchannel network was fabricated using a patterned negative photoresist on a silicon wafer using standard photolithography techniques. Cylindrical chamber molds were machined from polymethyl methacrylate (PMMA) using computer numerical control (CNC) milling. The final molds were then cast with polydimethylsiloxane (PDMS, Sylgard 184, Dow Corning, MI, USA) and plasma bonded. The localized heating zones were generated by using thermoelectric modules coupled with temperature controllers, and mounted on a three-dimensional printed platform. (**Supplementary Note 2**).

### 2.4. Numerical simulations

Bacterial transport into a microchamber, with a diameter of 7.6 mm and a height of 4 mm, was simulated in COMSOL Multiphysics (v6.0) under a steady laminar incompressible flow of a water-like fluid flow (with viscosity of 1 mPa.s and density of 1000 kg m^−3^), with bacteria modeled as 1 µm spherical particles subjected to Stokes drag and tracked using Lagrangian particle tracking. Residence times extracted from particle trajectories (**Supplementary Note 3**).

### 2.5 Temperature calibration

The surface temperature of the heating modules was visualized by an infrared thermal camera (FLIR Systems, OR, USA). To calibrate the set temperature on the controller and the temperature inside the chamber, we measured the internal temperature using a thermocouple (Thermo Fisher Scientific, MA, USA).

### 2.6 Device operation

A culture of *L. rhamnosus,* which was cultivated for 14–16 h, was diluted to 10^8^ CFU mL^−1^ with fresh medium. The microfluidic device was pre-filled with medium and then inoculated with 20 µL of the diluted culture. The device was left undisturbed for 30 min to allow cells to settle to the microchamber floor. Meanwhile, the heating zone controllers, which are separate from the device, were turned on and allowed to reach thermal equilibrium. The chambers were aligned with the heating zones and the fluid flow was controlled by a peristaltic pump (ISMATEC, Wertheim, Germany). Air bubbles from the medium stream were continuously removed by a bubble trap (PreciGenome, CA, USA) before the flow entered the device.

### 2.7 On-chip growth monitoring and analysis

Growth monitoring was done using a digital camera (NexiGo, OR, USA) integrated with a custom MATLAB script to capture sequential images of the chip every 15 min. Grayscale images were processed for visualization by converting pixel intensities to a colorized scale. Mean pixel intensities (I) within each microchamber were quantified over time and normalized to the maximum possible grayscale value 255. To enhance data quality, the intensity profiles were smoothed using a moving average filter with a 2 h window. We used a 24 h interval imaging method for experiments longer than 2 d. Metabolic activity was monitored daily by measuring the pH of the effluent using a pH meter (AQUASEARCHER, OHAUS, NJ, USA)

### 2.8 Statistical analysis

All data are presented as mean ± standard deviation from three independent measurements (n = 3 per condition). Statistical differences among multiple groups were evaluated using one-way analysis of variance (ANOVA). When a significant effect was detected, post hoc pairwise comparisons were performed using Tukey’s honest significant difference (HSD) test to control for multiple comparisons. Statistical significance was defined as p < 0.05. Reproducibility was assessed by calculating the coefficient of variation (CV = Standard Deviation between replicates/ Mean × 100) of CFU counts obtained from chamber S_1_ across independent replicates. All analyses were performed in MATLAB.

## 3 Results and discussion

### 3.1 Conceptual framework and design

The adaptive response of bacteria was found to be highly dependent on the spatial arrangement of the stressor; steep gradient (Zoheir et al., 2021) and smooth gradient (Nguyen et al., 2024). Motivated by this, we investigated how thermal gradient configuration influences bacterial growth dynamics. Our thermal adaptation platform operates by sustaining continuous growth and direct transport of cells from a low-temperature environment (T_1_) into a thermally stressful environment (T_2_) via controlled fluid flow (**Fig. 1a**).

**Figure 1.**
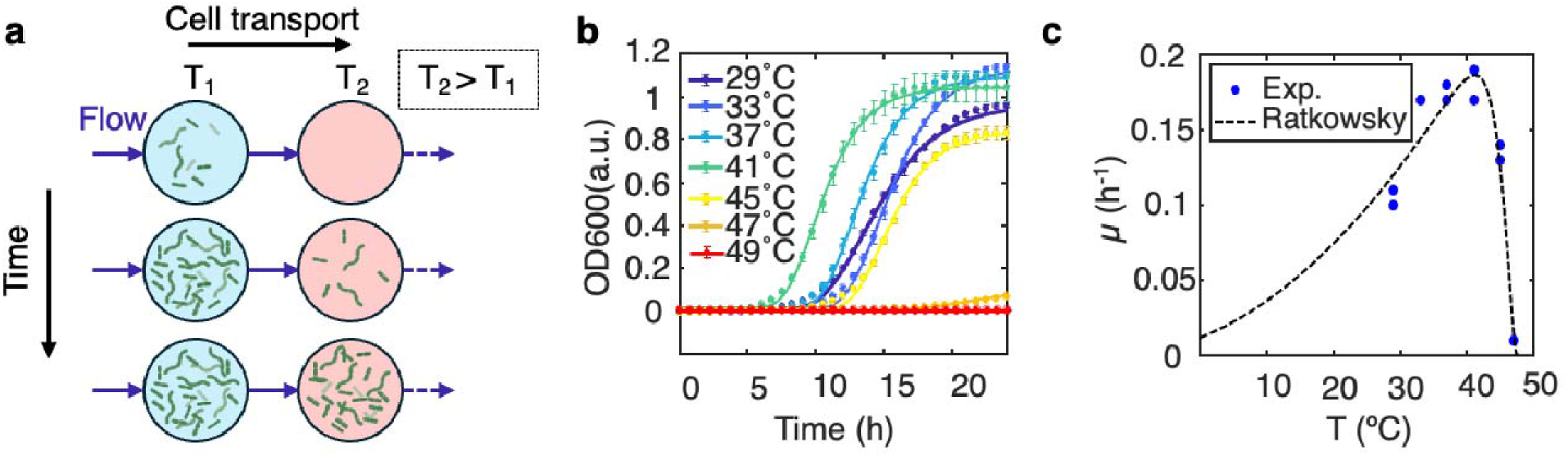
Conceptual design of the flow-based stepwise thermal adaptation of bacteria. a Schematic of the microfluidic architecture enabling stepwise exposure of bacterial populations to increasing thermal conditions. Cells are serially transported from low-temperature (T_1_) to high-temperature (T_2_) microchambers via precisely regulated medium flow. b Growth curves of *L. rhamnosus* at different temperatures to obtain T_1_ and T_2_. c. Effect of incubation temperature (T) on the maximum specific growth rate (µ_max_).

First, to identify appropriate values for T_1_ and T_2_ for this strain, we first quantified growth across a range of incubation temperatures (29–49 °C) using microwell plates and monitoring the optical density with inoculum size of 10^7^ CFU mL^−1^ (**Fig. 1b and Supplementary Note 1**). The optimal growth temperature for *L. rhamnosus* is 41 °C with a maximum specific growth rate (µ_max_) of 0.18 ± 0.01 h^−1^; at temperatures exceeding this, the growth rate decreased sharply. (**Fig. 1c**). The thermal response curve fits well to the Ratkowsky model(Ratkowsky et al., 1982) (**Supplementary Note 1**). The lag phase duration was 8 h at 41 °C and increased to 13 h at 45 °C (**Fig. S1**). The net growth yield was reduced by 20% at 45 °C compared to the optimum temperature. No growth was observed at 47 °C. Because the bacterial thermal response depends on the inoculation size(Augustin Jean-Christophe et al., 2000; Robinson et al., 2001), we repeated the experiments with a 10-fold diluted inoculum (∼10^8^ CFU mL^−1^) and found that cells could grow at 47 °C, however, no growth was observed at 49 °C. Additionally, increasing the inoculum size consistently shortened the lag phase by nearly 50% across all tested temperatures (29–45 °C), but no noticeable change was observed in net growth yield (maximum optical density of culture) (**Fig. S2**). CFU count comparison, while slightly decreased, showed evidence of growth and that incubation at 49 °C did not kill the cells (**Fig. S3**). Thus, 49 °C represents a sublethal thermal-stress condition, regardless of the inoculum size, for *L. rhamnosus*: proliferation ceases at this temperature, but cellular viability is preserved.

To generate segregated yet connected microenvironments, we used circular micro-bioreactors on a microfluidic device arranged on a temperature-controlled heating platform (**Fig. 2a and b**). Microchambers have a volume of 180 µL and were manufactured from polydimethylsiloxane (PDMS) (**inset, Fig. 2a**). Each chamber is then aligned on a millimeter-sized heating module where the temperature is controlled (**Fig. 2b**). Connections between the microchambers and nutrient delivery are provided by a network of microchannels (200 µm wide, 50 µm high) (**Fig. 2c, Supplementary Note 2**). The design ensures identical hydrodynamic conditions across all chambers (constant medium inflow), allowing temperature to remain the primary variable. To avoid biases arising from high cell density (Augustin Jean-Christophe et al., 2000), which can interfere with cellular adaptation, we incorporated a continuous serial dilution (CSD) unit that dilutes both the cells and their carried metabolites in the effluent stream before it enters the next chamber (**Fig. 2d**). In the CSD unit, 90% of the effluent from chamber S_i–1_ is discarded, and the remaining fraction is supplemented with fresh medium to re-establish a flow rate equivalent to that supplied to chamber S_i_.

**Figure 2.**
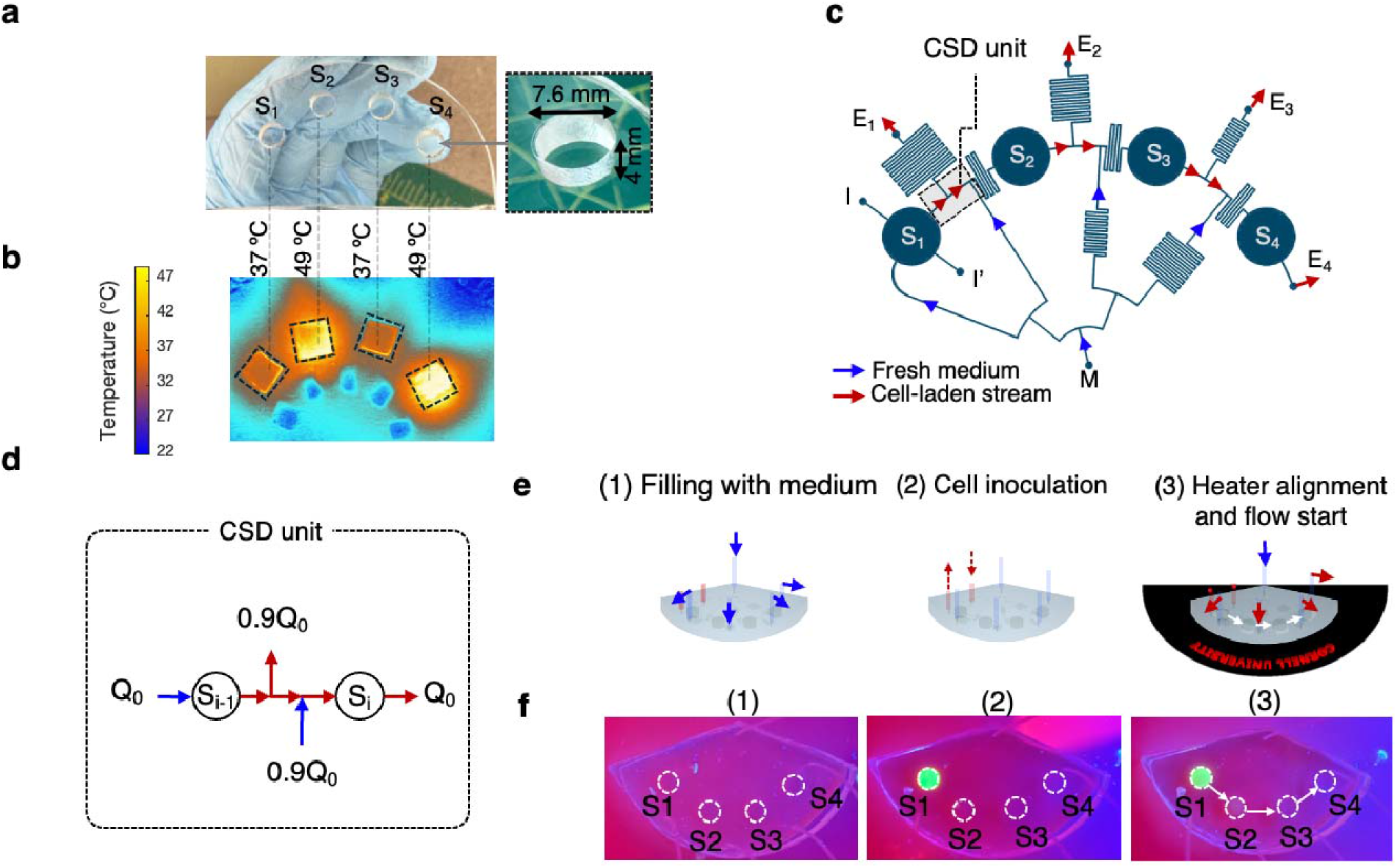
Design of the microfluidic platform for on-chip cultivation of bacteria on a thermal stress landscape. (a–b) Photograph of the microfluidic chamber array alongside the corresponding heater-generated temperature profile, illustrating sharp spatial confinement of thermal zones. c. Design of the microfluidic device illustrating the main structural components and flow pathway. d. Schematic of continuous serial dilution (CSD) unit. e. Operation steps of the microfluidic device. f. Localized inoculation demonstrated using fluorescein, highlighting precise delivery of cells into S_1_.

Operation of the microfluidic device consists of three steps: 1) filling the device with the growth medium, 2) inoculation of the cells to the device, and 3) aligning the microchambers on the heaters and starting the medium flow (**Fig. 2e**). Inoculation of cells to the system must occur in the first microchamber because the device is designed for unidirectional flow and serial dilution, requiring a defined starting point for controlled downstream propagation of cells. To validate the inoculation step, we filled the microfluidic chip with water and introduced a fluorescein solution into chamber S_1_. Fluorescence imaging showed the highest signal in S_1_, confirming precise inoculation (**Fig. 2e**). After 4 h of water flow at 576 µL h^−1^, only a weak signal appeared in S_2_, demonstrating proper serial connectivity and effective operation of the serial dilution unit (**Fig. 2f, step 3**).

### 3.2 Experimental setup

The experimental setup consists of three main components: 1) a microfluidic device and flow control, 2) a heating module for generating localized temperatures, and 3) an imaging module (**Fig. 3a**). Continuous medium flow is provided by a peristaltic pump supplementing the cultures growing in the microchambers. Air bubbles in the incoming medium are continuously removed by a bubble trap to prevent flow imbalance in the fluidic module. Heat is supplied to microchambers by four thermoelectric modules (Drebushchak, 2008) coupled with controllers for temperature adjustment (**Fig. 3b** and **Fig. S6** for the electrical circuit) all mounted on a custom three-dimensional printed mount. The voltage input to the thermoelectric modules is adjusted by a power supply. For consistency and efficiency of the heat pump, we cooled down the lower part of the module using a cooling fan. Heat transfer was maximized by filling the space between heater and temperature sensor (thermistor) using a thermally conductive paste (**Fig. 3b**). The thickness of the PDMS layer of the floor of the chambers is ∼2 mm to maximize the efficiency of heat transfer from heater surface to the medium inside microchambers. A photograph of the assembled setup is shown in **Fig. 3c**.

**Figure 3.**
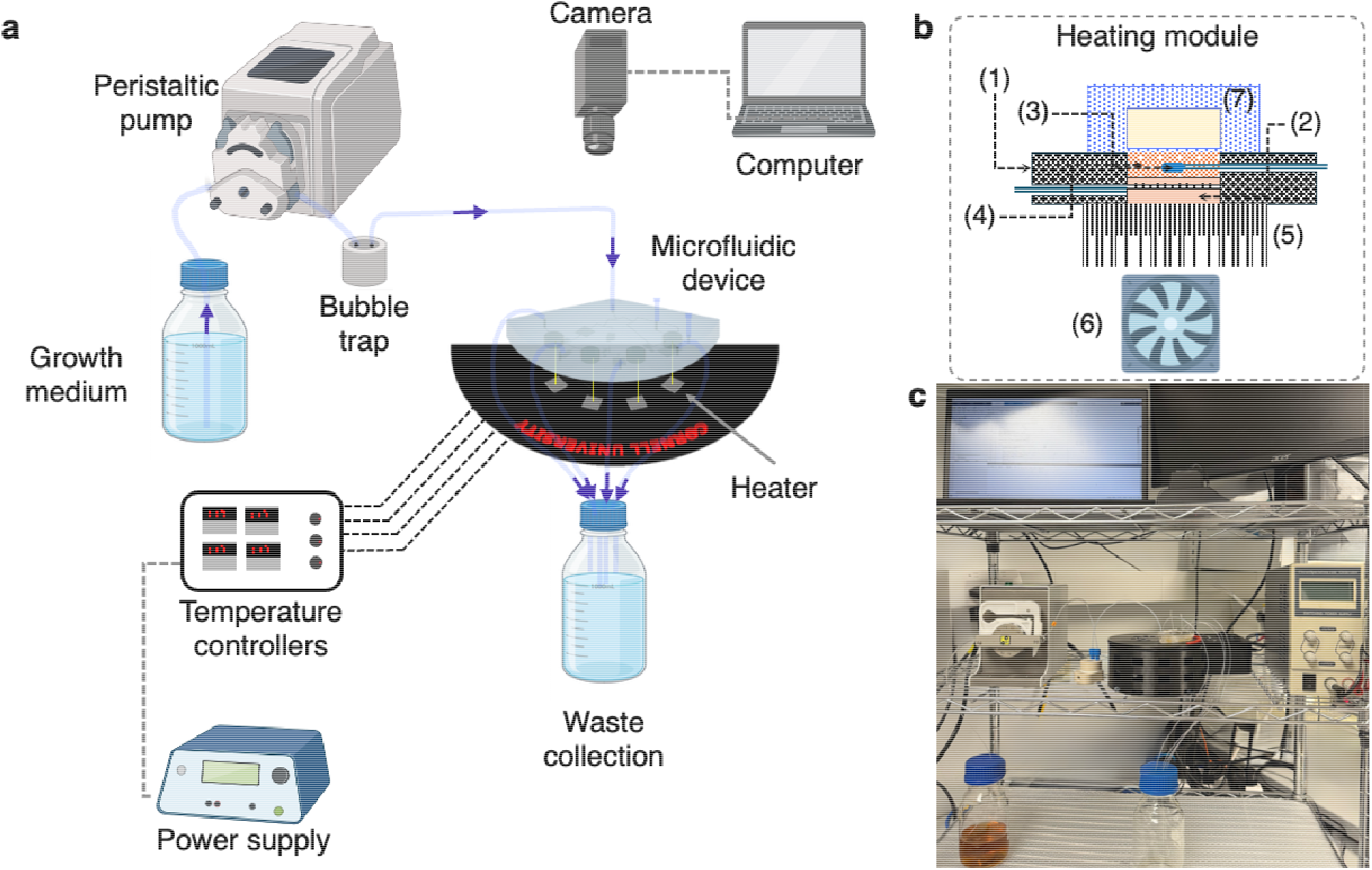
Experimental platform. a. Schematic of the on-chip thermal adaptation setup. b. Components of the heating module: (1) 3D-printed mount, (2) thermoelectric module, (3) thermally conductive filler, (4) temperature sensor, (5) heat sink, (6) cooling fan, and (7) PDMS microchamber. c. Photograph of the assembled setup.

### 3.3 Platform characterization

Two main components of our microfluidic system, namely thermal and hydrodynamic conditions, were investigated separately. Each microchamber is a bioreactor heated from below; the heat is supplied to the growing biomass inside the chambers through a proportional controller where the heater surface temperature (T_H_) is maintained (**Fig. 4a**). We used proportional controllers because they integrate easily with our system and provide stable regulation of T_H_ around the desired setpoint (**Fig. S6**). While proportional control does not completely eliminate steady-state fluctuations, it does maintain the temperature within a narrow range suitable for our experiments. Consequently, T_H_ exhibited only slight fluctuations during operation (**Fig. S8**). We developed a mathematical model to predict the temperature variation within the liquid inside the chamber. Experimentally we found a small thermal resistance attributed to the PDMS layer on the chamber ceiling which generated a 0.1 °C temperature fluctuation (**Supplementary Note 3**). Additionally, there is a heat loss within the chamber which changes the final temperature of the growth medium (T_M_). To account for this, we calibrated each individual chamber/heater module for each temperature zone by using an external digital thermometer placed inside the chamber (**Fig. 4b**). We did not include the effect of fluid flow. Although all heaters had linear T_H_–T_M_ correlation, the slopes were different which captures the overall resistance between the heater and the liquid inside the microchambers, and the slight inherent differences between the thermoelectric modules.

**Figure 4.**
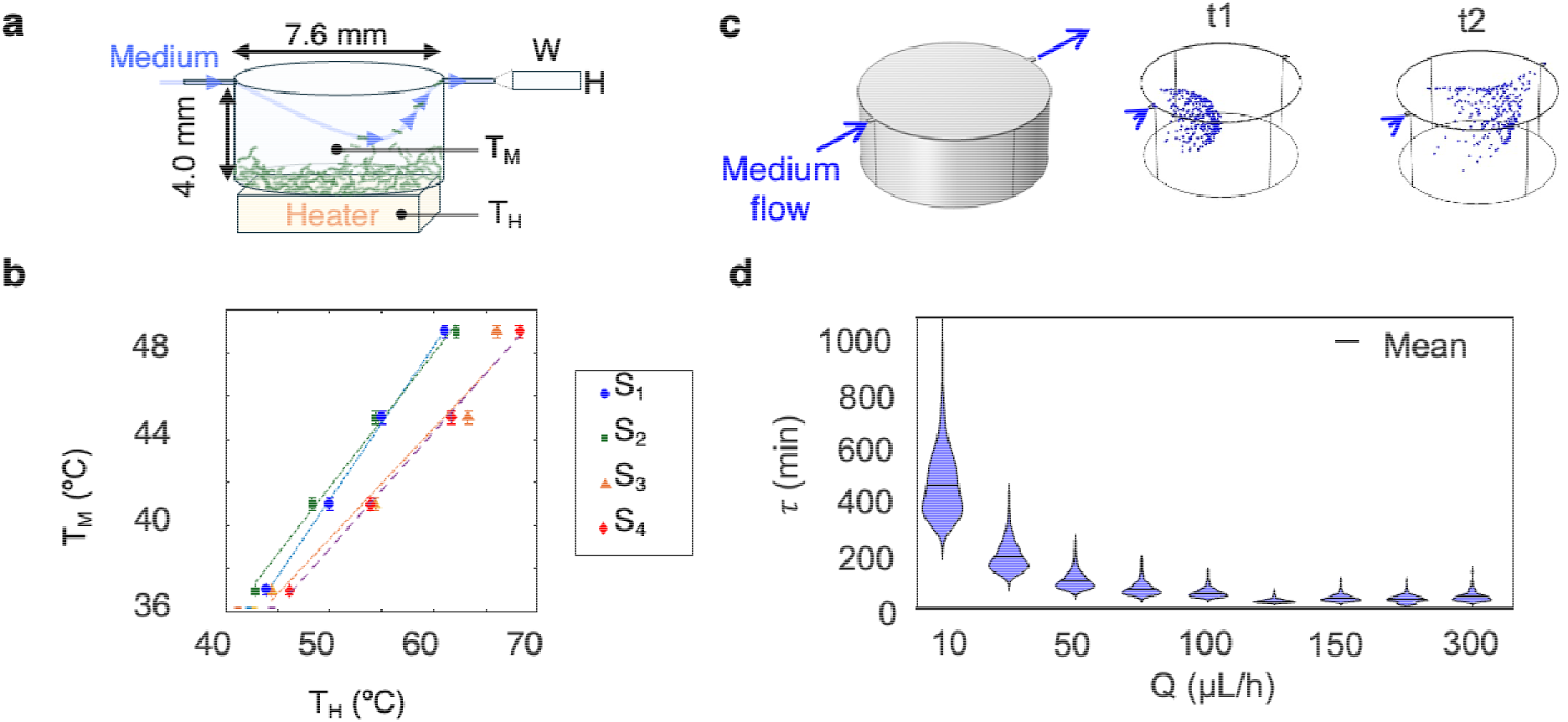
Characterizing the microfluidic device and heating modules. (a) Schematic showing the micro-bioreactor. The heater generates a surface temperature (T_H_) which increases the temperature of the medium inside the chamber (T_M_). (b) Calibration of T_H_ and T_M_ within the chamber for each heater set. (c) Geometry of the microchamber used for simulating cell transport within the chamber and the visual distribution of 1-µm particles initially released at inlet at time t1 (flow expansion) and t2 (flow contraction) where they exit the chamber. Flow rate is 100 µL h^−1^. (d) Dependence of cells residence time in the chamber during the transport.

During the experiment, single cells are predominantly transported by flow convection to the next in-line microchamber as controlled by inlet fluid flow to each chamber. This altered fluid flow impacts the residence time of cells in the chambers. To account for this, we used particle tracking simulations to predict the residence time of bacteria in the chambers (see **Fig. 4c** and **Supplementary Note 3**). The residence time decreased exponentially with every incremental increase in flow rate. (**Fig. 4d**).

The minimum input flow rate (Q_min_) to the microchambers was estimated by considering the maximum specific growth rate ( µ_max_) of *L.rhamnosus,* such that Q_min_ = µ_max_, assuming a well-mixed chemostat (Gresham and Dunham, 2014), that allows the cells to grow within the chamber before exiting the chamber. The maximum specific growth rate was observed to be 0.2 h^−1^ at 37 °C (**Fig. 1c**). Assuming a well-mixed bioreactor, the required flow rate (Q) is the volume of the reactor multiplied by the growth rate (Q = 180 µL ྾ 0.2 h^−1^ = 36 µL h^−1^). The particle trajectory simulations showed that a flow rate of 36 µL h^−1^ established a particle residence time of 3.3 h in each chamber (**Fig. 4d**), which is comparable to the theoretical doubling time (t_d_= ln(2)/µ_max_= 3.5 h). However, experimental testing of two flow rates (36 and 144 µL h^−1^) showed that the minimum flow rate is insufficient for long-term cultivation and higher flow rate enhanced biomass growth and cell transport by delivering more nutrients and washing away metabolic waste (**Fig. S9**). Therefore, we used a flow rate of 144 µL h^−1^ to ensure the efficient nutrient delivery and cell transport.

### 3.4 Growth under no thermal stress

To evaluate bacterial growth in the system and verify reproducibility, a control experiment was done where all microchambers were maintained at a constant 37 °C (**Fig. 5a**). The growth of *L. rhamnosus* generated a white-colored biomass such that the pixel intensity of the appearance of the biomass could be tracked over time (**Fig. 5b** and **Fig. S10**). The microchambers were labelled S_1_ through to S_4_ and the flow within the platform carries the cells in series from S1 to each subsequent chamber. The growth dynamics in microchambers S_1_ through S_4_ indicated a gradual growth initiation in each chamber, with S_1_ showing growth within 5 h (**Fig. 5c**). The absence of visible growth inside chamber S_4_ during the course of the 48 h control experiment is likely due to the low concentration of cells reaching that chamber. The use of CSD units (**Fig. 2d and Fig. S13**) effectively diluted the cell population, reducing the initial cell density and thereby prolonging the lag phase before growth resumed. The extension of lag time in microchambers S_1_ through S_3_ and no growth in S_4_, is consistent with the inoculum size-dependence in off-chip experiments (**Fig. S2**).

**Figure 5.**
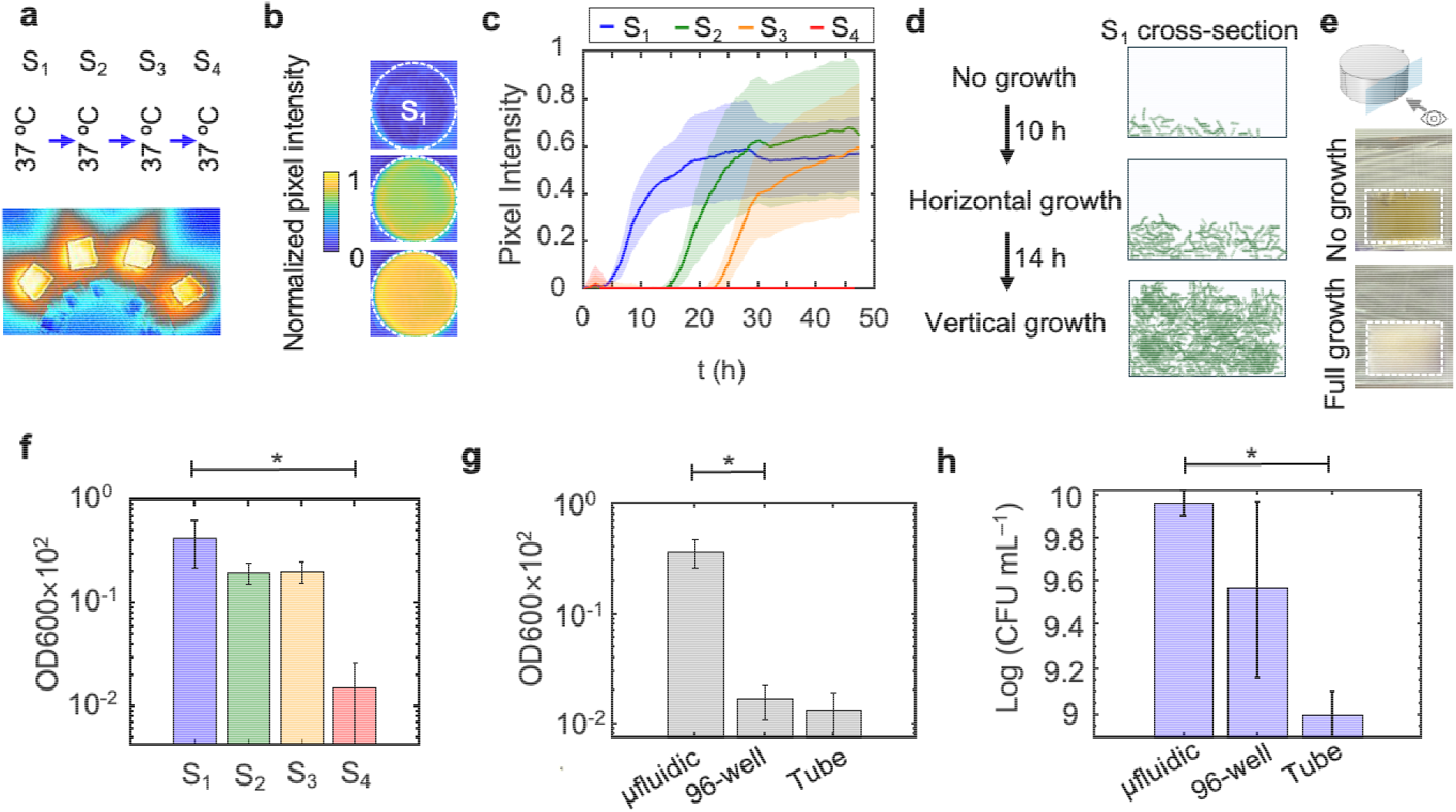
Growth of *L. rhamnosus* under no stress condition. On-chip growth control at 37 °C. (a) Infrared image of the thermal stand showing heated spots are maintained at 37 °C. Dark blue color indicates 22 °C and yellow indicates 37 °C. (b) Representative images showing pixel intensity contrast between an empty chamber and one filled with *L. rhamnosus* biomass. (c) Temporal evolution of chamber pixel intensity profiles (mean ± s.d., *n* = 3 replicates). (d) Schematic of the growth stages within a microchamber. (e) Photographs of the microchamber side view for no growth and full growth. (f) Optical density (OD_600_) of 100×-diluted cultures extracted from S_1_–S_4_ after incubation. Error bars represent standard deviation of three replicates. (g) Comparison of OD_600_ and colony-forming unit (CFU) concentrations obtained from microfluidic cultures, 96-well plate cultures, and test-tube cultures. Data are shown as individual replicates with mean ± SD (n = 3 independent measurements per condition). Statistical differences among groups were assessed using one-way ANOVA followed by Tukey’s HSD post hoc test. Significant differences are indicated by an asterisk (*p < 0.001).

Bacterial proliferation within each chamber consists of three main stages: 1) no growth, or lag phase, 2) horizontal growth, when cells start to spread on the chamber floor, and 3) vertical growth (**Fig. 5d** and **Supplementary Note 4**). The side view of the chamber S_1_ after 48 h of incubation shows an opaque biomass filled the chamber (**Fig. 5e**). After 48 h, optical density measurements of cultures extracted from each chamber revealed differences in cell proliferation in each chamber. S_1_ exhibited the highest cell density and S_4_ the lowest (**Fig. 5f**). Compared with conventional batch cultures at 37 °C, the microfluidic chambers produced ∼10-fold higher optical density than 96-well plates and test-tube cultures (**Fig. 5g**). Cells recovered from the chambers remained viable, reaching 10^10^ CFU mL^−1^ (**Fig. 5h**), which is ∼10-fold higher than that of test-tube cultures. The small variation among replicates of CFU measurements in S_1_ (coefficient of variation = 0.6%) indicates consistent and reproducible performance of the microfluidic system.

### 3.5 Influence of thermal stress landscape

When we exposed the cells to a uniform thermal stress landscape by maintaining all microchambers at 49 °C, no growth was observed after 3 days of cultivation (**Fig. 6a**). We postulated that the continuous fluid flow and reduced growth rate at the sub-optimal temperature prevented the establishment of growth within chambers. To test this, we maintained S_1_ at 37 °C so that the bacteria could proliferate in this chamber and kept the rest of the chambers at 49 °C. Bacterial growth filled the first chamber after 2 days: However, no visible biomass was observed in subsequent chambers that were held at 49 °C (**Fig. 6b**).

**Figure 6.**
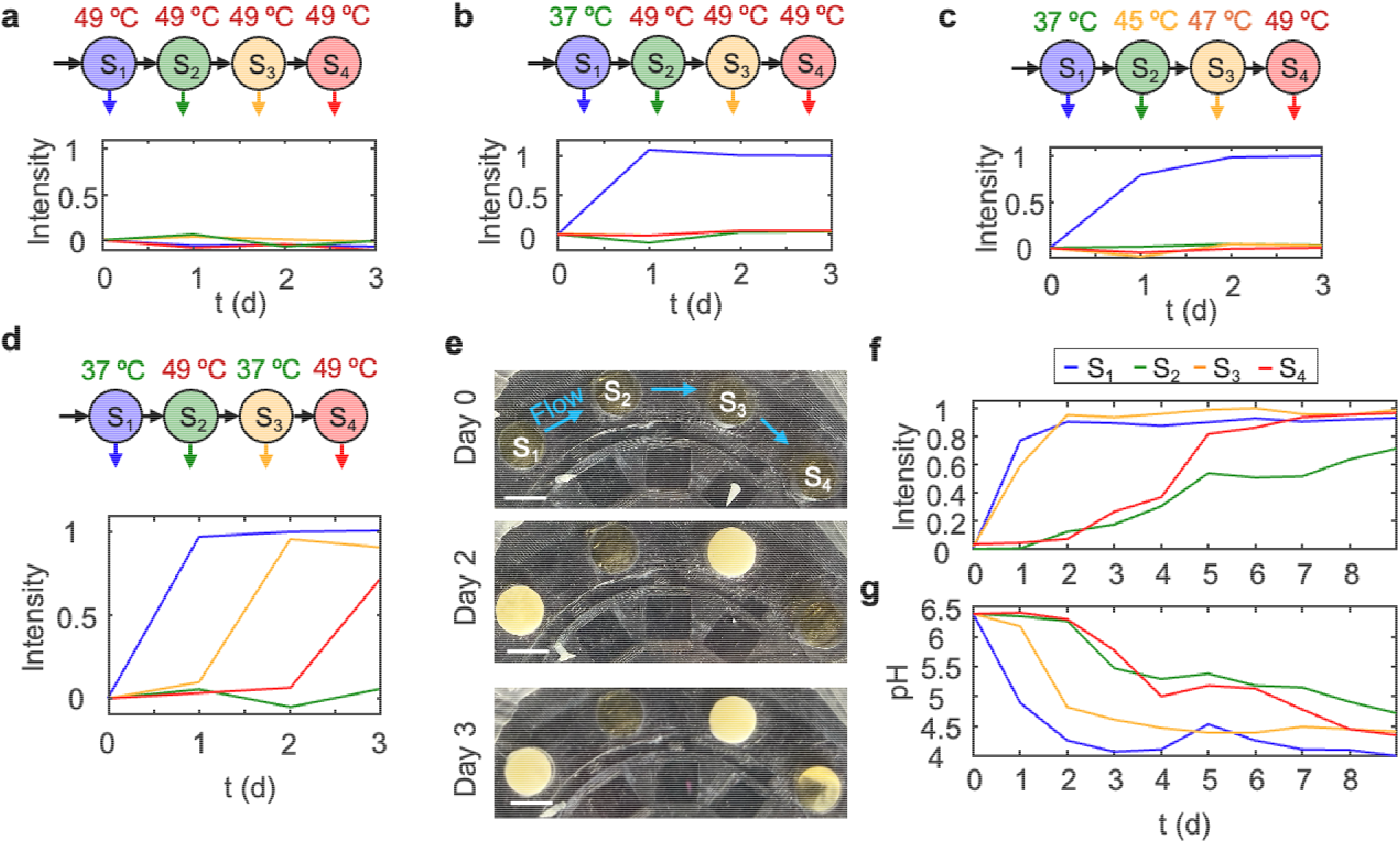
(a-d) influence of different thermal landscape configuration on the growth and adaptation of *L. rhamnosus*. (e) Images of the chambers showing biomass formation in S_4_ chamber after 3 d of incubation. Scale bars: 1 cm. (f) Average chamber pixel intensity profiles during long-term incubation under periodic thermal stress, representing the spatial distribution and evolution of biomass over time. (g) Decrease in effluent pH from each chamber, indicating metabolic activity associated with biomass growth on-chip.

Previously gradients of stressors have been effective in producing stressor in producing adaptations toward oxidative stress (Nguyen et al., 2024) and antibiotics (Zoheir et al., 2021). Therefore, we designed a thermal gradient (37–49 °C) informed by the off-chip thermal response curve (**Fig. 6c**). However, no visible biomass was observed after 3 days of incubation in the subsequent chambers (**Fig. 6c**).

Thermal adaptation of bacteria has been reported when alternating temporal temperature change (Mongold et al., 1996; Bennett and Lenski, 1997). We designed a periodic thermal stress landscape (S_1_ = 37 °C, S_2_ = 49 °C, S_3_ = 37 °C, S_4_ = 49 °C) (**Fig. 6d and e**). In this experiment, after 3 days we observed biomass in S1 and S3 and in S4. Longer incubation for 9 days revealed that the biomass was observed in both S_2_ and S_4_, however, the accumulation and generation was continuously higher in S_4_ **(Fig. 6f**). The long lag time (48 h) in S_2_ and S_4_ originates from thermal stress since it is much longer than the lag time originating from serial dilution (15 h) (**Fig. 5c**).

To assess the metabolic activity of cells during the on-chip cultivation, we measured the pH of the outlet effluents from each chamber (E_1_–E_4_ in **Fig.2c**). The decrease in pH is an indication of the organic acids produced by *L. rhamnosus* providing evidence of an active culture in each chamber (Fayol-Messaoudi Domitille et al., 2005) (**Fig. 6g**). When chambers S_1_ and S_3_ reached their maximum cell capacity (see **Fig. 5e**), the pH reached a plateau at ∼4.3 indicating that the cells are actively dividing and metabolizing despite the chamber having reached maximum cell density. The culture is maintained at its late exponential phase throughout the experiment due to the continuous refreshment of nutrients and waste removal, which resembles a chemostat (Pirt, 1975). The 48 h-lag time for S_2_ and S_4_ obtained from chambers intensity measurements closely matches those of the pH profile. Bacteria grew much slower in S_2_ and S_4_ compared to S_1_ and S_3_ due to the thermal stress, consistent with the off-chip growth kinetics (**Fig. 1c**). After 9 days of cultivation, the extracted cultures from chambers had comparable optical densities showing the growth of bacteria in all chambers (**Table S3**).

To assess the lasting impact of thermal stress, we evaluated bacterial adaptation using an off chip thermal response assay (see methods). No growth was observed in a large scale, off chip culture carried out at 49 °C after 24 h (**Fig. S14**), suggesting that acclimation (Bennett and Lenski, 1997) or phenotypic plasticity (Hurtado-Bautista et al., 2021) may enable the bacteria to proliferate on chip at high temperatures while remaining incapable of growth under identical off chip conditions. A similar phenomenon has been observed in *Escherichia coli*, which could grow at high antibiotic concentrations on a microfluidic chip but not in conventional off chip cultures (Zoheir et al., 2021).

*L. rhamnosus* exhibits a chaining phenotype arising from incomplete cell division (Claes et al., 2012). We further postulate that the distinctive chaining behavior of *L. rhamnosus* (**Fig. S15**) promotes the formation of tightly coiled cellular aggregates (**Fig. S16**). We observed that high shear within the microchannel disrupts cell chains during transport between wells (**Fig. S16**)(Gomand et al., 2020). This fragmentation increases the number of single cells entering the high-thermal-stress chamber, which may enhance overall survival and subsequent growth.

## 4 Conclusions

Temperature is a crucial factor in determining the viability, growth, and proliferation of bacteria, including probiotics which are thermally stressed during industrial processing and storage. In this paper, we developed an integrated microfluidic platform to assess the thermal response of *L. rhamnosus* to different thermal stress configurations. When exposed to a landscape of near optimal temperature (all chambers at 37 °C), we observed growth in all chambers within 48 h and the extracted cells from the microchambers were viable. When setting all chambers at the thermal stress condition (49 °C), no growth was observed even after 3 days. However, in a periodic thermally stressed condition (37 °C, 49 °C, 37 °C, 49 °C), there was an extension of growth lag and we observed growth after 5 days of incubation in all chambers including those held at 49 °C. We also monitored the metabolic activity for 9 days by tracking the pH of each chamber through effluent analysis.

In summary, using an integrated microfluidic platform, we demonstrated that the spatial profile of thermal gradients within a microfluidic environment plays a role in shaping bacterial growth dynamics and adaptive responses. The platform enables continuous sampling from individual chambers, delivers reproducible culturing conditions, and supports automation, making it well-suited for long-term adaptive laboratory evolution (ALE) experiments. These findings indicate that microscale environmental structure can influence microbial proliferation and adaptation not only through temperature itself, but also by modulating cell transport, spatial exposure to stress, and population heterogeneity across compartments. In the context of probiotics, such control over microenvironmental conditions may be leveraged to improve functional performance by enabling selection or maintenance of more stress-tolerant and metabolically active subpopulations, increasing viable cell delivery to the host, and enhancing robustness during manufacturing, storage, and gastrointestinal transit. Future work will focus on implementing ALE experiments on this platform, incorporating multiple stressors, such as combining chemical and thermal stressors in one experiment, and extending the approach to a broader range of bacterial species.

## Author contributions

A. K. performed conceptualization, investigation, methodology, formal analysis, wrote the original draft, wrote, reviewed and edited the draft. M.Y. performed methodology, reviewed and edited the draft. S.Z. performed methodology, formal analysis, wrote, reviewed and edited the draft. G.P.A., G.F., and P.M. performed conceptualization and wrote, reviewed and edited the draft. K.W.S. provided resources, and wrote, reviewed and edited the draft. A.A. performed conceptualization, funding acquisition, project administration, supervision, writing – editing and reviewing and provided resources.

## Supporting information

Supporting Information and Discussion

## Acknowledgement

We thank Dr. Kelley Donaghy for her assistance in editing the manuscript. This research was performed under contract no. W911QY2210007 with the DEVCOM Soldier Center, Soldier Sustainment Directorate of the United States Army, and published under public release approval number PR2026-3299. This work was performed in part at the Cornell NanoScale Facility, a member of the National Nanotechnology Coordinated Infrastructure (NNCI), which is supported by the National Science Foundation (Grant NNCI-2025233).

## Supplementary Information

Raw data that supports the figures in this manuscript can be found at: https://doi.org/10.5281/zenodo.15091594

## Generative Artificial Intelligence (AI) Statement

The authors certify that generative AI was not used in preparing this article. Non-generative AI, such as spelling and grammar checkers in Office 365 and Google Docs, and citation management software, was used. All instances when non-generative AI was used were reviewed by the authors and editors.

