## Supporting Information and Discussion for "Microfluidic cultivation enables probiotic growth under thermal stress"

**Supplementary Tables Legend**

**Table S1.** Colony forming units (CFU) density for different dilution factors

**Table S2.** Parameters and values considered for mathematical modeling of heat transfer to microchamber

**Table S3.** Optical density (OD600) of cultures extracted from the experiment on the thermal stress landscape.

**Supplementary Figures Legend**

**Fig. S1**. Thermal response of wild-type *L. rhamnosus* at different incubation temperatures

**Fig. S2**. Effect of inoculum on thermal response of *L. rhamnosus*.

**Fig. S3**. Cell viability after incubation at 49°C for 18 h.

**Fig. S4**. Microfluidic network design.

**Fig. S5**. Fabrication of the multiscale microfluidic device.

**Fig. S6**. Electrical design of µ-Therm system

**Fig. S7**. Schematic representation of the heat transfer model for the microchamber system.

**Fig. S8**. Comparing oscillation of temperature on the µ-Therm heater T_h_ and inside the microchamber T_L_

**Fig. S9**. Effect of refreshing medium flow rate on bacterial growth in the microfluidic device.

**Fig. S10**. Intensity profile of a microchamber over the cultivation time.

**Fig. S11**. Schematic showing the phases of bacterial growth in the microchamber

**Fig. S12**. Numerical simulation and experimental intensity profile of biomass settlement in a microchamber.

**Fig. S13**. Continuous serial dilution (CSD) unit operation between chambers S_1_ and S_2_.

**Fig. S14**. Growth curves at 49 ºC of the on-chip grown cultures in the periodic thermal landscape.

**Fig. S15**. Phase-contrast microscopy images of cells collected from the S1 chamber after 48 h of incubation

**Fig. S16**. False-colored image of the CSD unit showing the transport of fluctuates between microchambers.

**Supplementary Notes Legend**

**Note 1.** Growth rate of L. rhamnosus at different temperatures

**Note 2.** Design and fabrication of microfluidic system

**Note 3.** Thermal and hydrodynamic characterization

**Note 4.** Biomass formation and dynamics in microchambers

**Table of contents for Supplementary Notes**

**Note 1. Growth rate of L. rhamnosus at different temperatures	4**

A. Culturing and growth monitoring	4

B. Optical density and cell density calibration	4

C. Growth curves and analysis	5

D. Influence of inoculum size	7

E. Viability of thermal-stressed cultures	8

**Note 2. Design and fabrication of microfluidic system	10**

A. Microfluidic channel network design	10

B. Microfabrication	11

C. Electrical circuit design for microheaters	12

**Note 3. Thermal and hydrodynamic characterization	14**

A. Mathematical modeling of heat transfer to microchambers	14

B. Residence time of cells in chambers	17

**Note 4. Biomass formation and dynamics in microchambers	19**

**References	24**

### Note 1. Growth rate of *L. rhamnosus* at different temperatures

#### A. Culturing and growth monitoring

A single colony of *L. rhamnosus* GG was picked from a MRS-agar plate, inoculated to 15 mL of MRS broth, and incubated at 37 °C with 180 rpm shaking overnight (Final OD600 = 1.35). A stock of 300 µL aliquots was prepared in glycerol (OD600 = 1.0) and stored at -80°C. We used a 96-well plate to prepare the 200 µL cultures (three replicates) by diluting the thawed frozen-cultures to a final OD600 of ~0.01 (~2×10^6^ CFU mL^–1^) by adding fresh MRS broth. The plate was then incubated at the specific temperatures in a plate reader (SpectraMax iD3 Multi-Mode) with medium shaking, and the OD600 of the wells was tracked over the time. Note that we used a fresh set of cultures for each temperature experiment. We obtained the growth parameters, including the maximum optical cell density (A), maximum specific growth rate (μ), and lag time (λ), by fitting the data to the reparametrized Gompertz model (Zwietering et al., 1990) (**Eq. 7**).

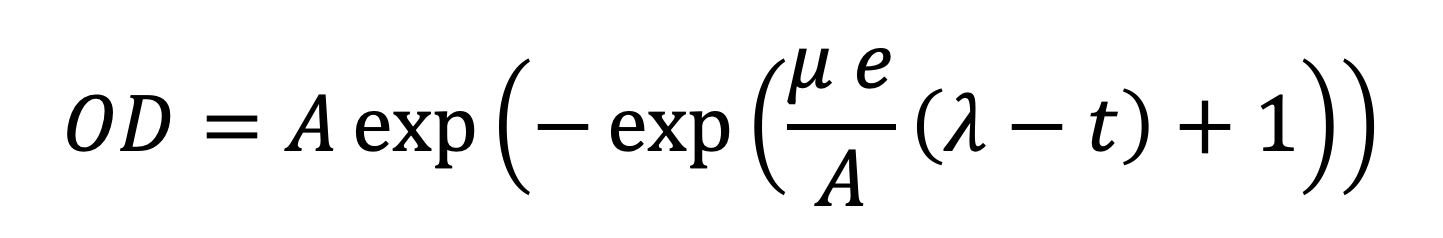
 **(1)**

where e ≈ 2.718 is the Euler number.

#### B. Optical density and cell density calibration

To ensure the consistency of the monitored OD600 with the actual cell density in the cultures, we calculated the colony forming units (CFUs). A 5 mL culture of *L. rhamnosus* was incubated for 16 h at 37 ºC in a test tube until it reached an optical density of 1.36. The culture was then serially diluted using phosphate buffer saline (PBS) and different dilutions (10^–5^, 10^–6^, and 10^–7^) were plated on agar and subsequently incubated at 37 ºC for 44 h until the appearance of colonies. Note that each dilution of culture had three replicates. The average cell density for the OD600 of 1.36 was ~10^9^ CFU mL–1 (**Table S1**). Therefore, 1 OD600 corresponds to ~10^9^ CFU mL^–1^.

**Table S1.** Colony forming units (CFU) density for different dilution factors (OD600 = 1.36).

| **Culture dilution factor** | **CFU mL^–1^** |
| --- | --- |
| 10^–5^ | 1×10^9^ |
| 10^–6^ | 2×10^8^ |
| 10^–7^ | 1×10^7^ |

#### C. Growth curves and analysis

Growth curves of *L. rhamnosus* at several temperatures, 29, 33, 37, 41, 45, 47, and 49 °C, are shown in **Fig. S5(a)**. Maximum growth rate μ_max_= 0.18 ± 0.01 h^–1^ was accomplished at 41 °C, therefore, the optimal growth temperature for this strain is close to T_opt_ = 41 °C (**Fig. S5b**). Note that for all experiments, the inoculum size was 10^8^ CFU mL^–1^. The alteration of growth rate under suboptimal conditions depends on whether the temperature lies below or above the optimum temperature (T_opt_). The rate of growth rate decay is steeper in supra-optimal temperatures. We fitted the growth rate data to the Ratkowsky square-root model(Ratkowsky et al., 1982) presented in Eq. **8**.

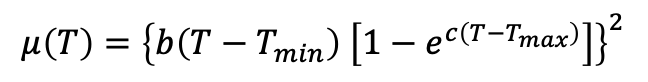
(2)

Where T_min_ and T_max_ are theoretical minimum and maximum, *b* is the slope parameter in the suboptimal (low–to–moderate) temperature range, and *c* is the curvature parameter describing how growth declines as temperature approaches T_max_. The values obtained from fitting are as follows: T_min_ = –13 °C, T_max_ = 47 °C, *b* = 0.008 $\sqrt{h^{-1}}/^{\circ}C$, and *c* = 0.54 °C^–1^. The calculated theoretical T_opt_ was 41°C. All predicted values were constrained to lie within biologically meaningful ranges, except for T_min_, which was constrained to exceed the freezing point of the medium (approximately 0°C). Growth was suppressed when temperature was elevated to 47 °C, above 49 °C growth ceased entirely.

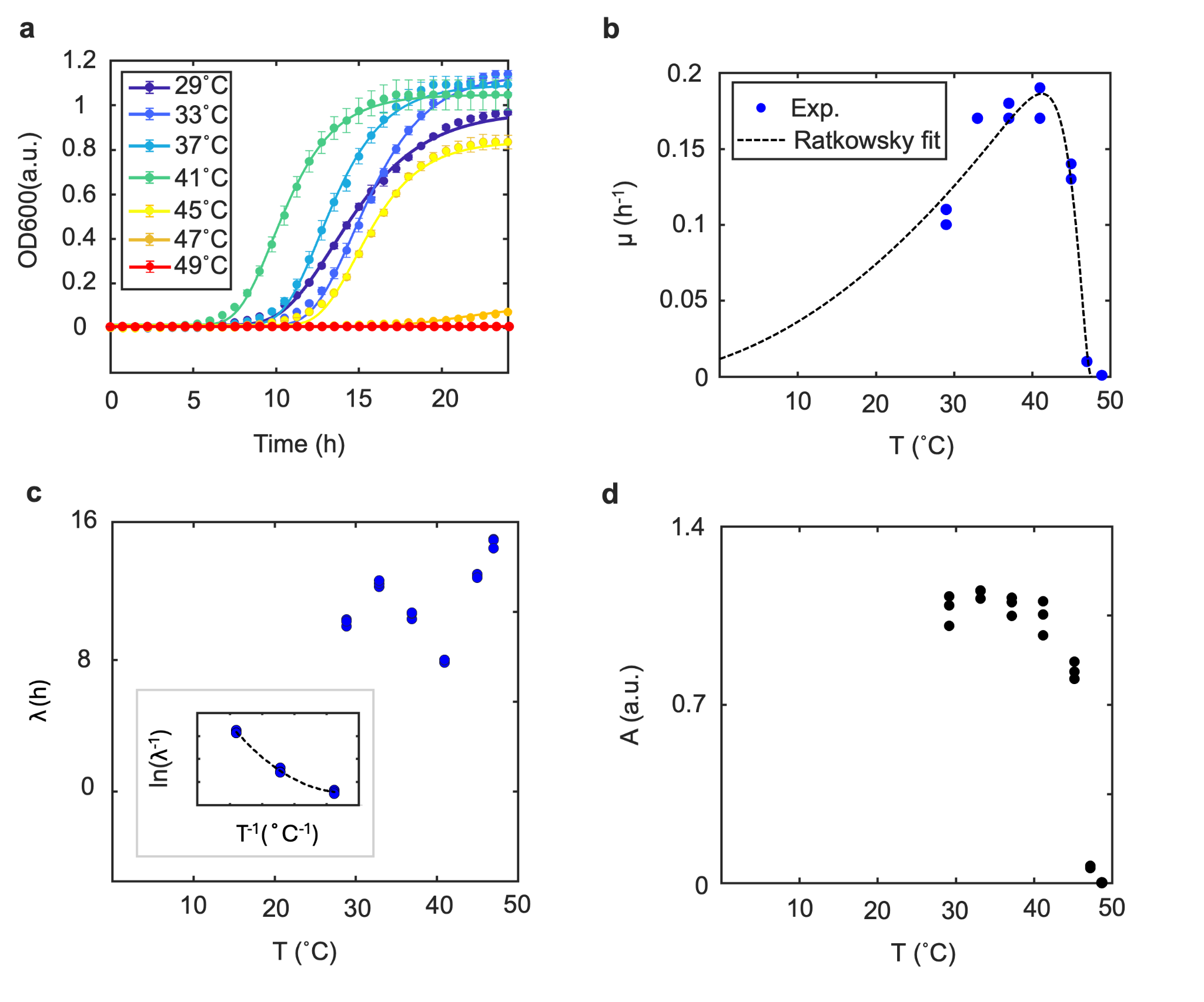

**Fig. S1** Thermal response of wild-type *L. rhamnosus* at different incubation temperatures. (**a)** Optical density at 600 nm (OD600) of cultures (N = 3 replicates). Solid lines indicate the fit to the reparameterized Gompertz model(Zwietering et al., 1990). (**b)** Maximum specific growth rate (µ) as a function of temperature. (**c)** Lag phase duration under different thermal conditions; inset shows 33–41 °C fitted to the model of (Daughtry et al., 1997). (**d)** Net growth yield (*A*) reduced under heat stress.

During the lag phase, the cell density remains practically stable. Temperature influenced the duration of lag phase duration λ of *L. rhamnosus* (**Fig. S5c**) depending on the temperature range. Lag time was prolonged when temperature increased from 29 °C to 33 °C, however, we observed a reversed trend until T_opt_. The lag time increased drastically beyond the optimal temperature, indicating a prolonged adaptation period of the cells under thermal stress. The data were subsequently well-fitted to the model proposed by Daughtry et al.(Daughtry et al., 1997) as presented in Eq. (**3**), using temperatures in the range of 33–41 °C.

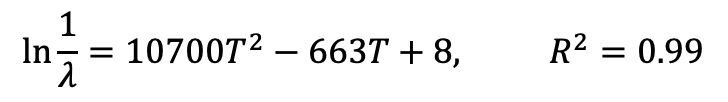
**(3)**

Maximum OD600 (*A*) indicates the maximum density of cells which occur at the stationary phase of bacterial growth. Suboptimal temperature incubations (29–41 °C) resulted in a final net growth of *A* = 1.1 ± 0.05 with minimal variation with temperature. However, elevation of temperature to 45 °C reduced the net growth to 0.8 ± 0.03 and increasing the temperature to 47 °C ceased all growth. Overall, the temperature for maximum growth rate of *L. rhamnosus* is 41 °C and temperatures higher than 45 °C are considered as thermal stress on cells.

#### D. Influence of inoculum size

Previous studies showed the dependence of inoculum size on the thermal response of various bacterial strains(Mackey and Kerridge, 1988; Augustin Jean-Christophe et al., 2000; Koutsoumanis and Sofos, 2005, 2005; De Silvestri et al., 2018) obtained the growth parameters for sub-cultures with a 10-fold concentrated inoculum (10X, ~10^7^ CFU/mL) and compared it to the original inoculums (1X, ~10^6^ CFU/mL) used in the previous section at different temperatures. Increasing inoculum size did not alter the overall trend of growth parameters between 29–45 °C; however, the μ_max_ was sevenfold higher compared to the smaller inoculum (**Fig. S7a**). No detectable growth was observed at 49 °C under either of conditions. The 10-fold higher initial cell density reduced the lag phase duration by approximately half (𝜆_10X_/𝜆_1X_ = 0.5) and was independent of temperature (**Fig. S7b**). The net growth was consistent between the two inoculum sizes with the expectation of 47 °C for which we obtained *A* = 0.4. We only observed the inoculum size effect when incubation temperature exceeds 47 °C where culture cannot initiate growth in low initial cell density but it grows well by increasing the inoculum size. Inoculum-size dependence of thermal response may result from cellular chemical interactions(Ardré et al., 2022), stochastic single-cell stress responses(Baranyi, 1998), and the higher probability of viable growth with larger inocula under low survival condition(Robinson et al., 2001). In summary, growth of *L. rhamnosus* is severely suppressed when temperature exceeds 49 °C, and the inoculum size influences the growth initiation at 47 °C.

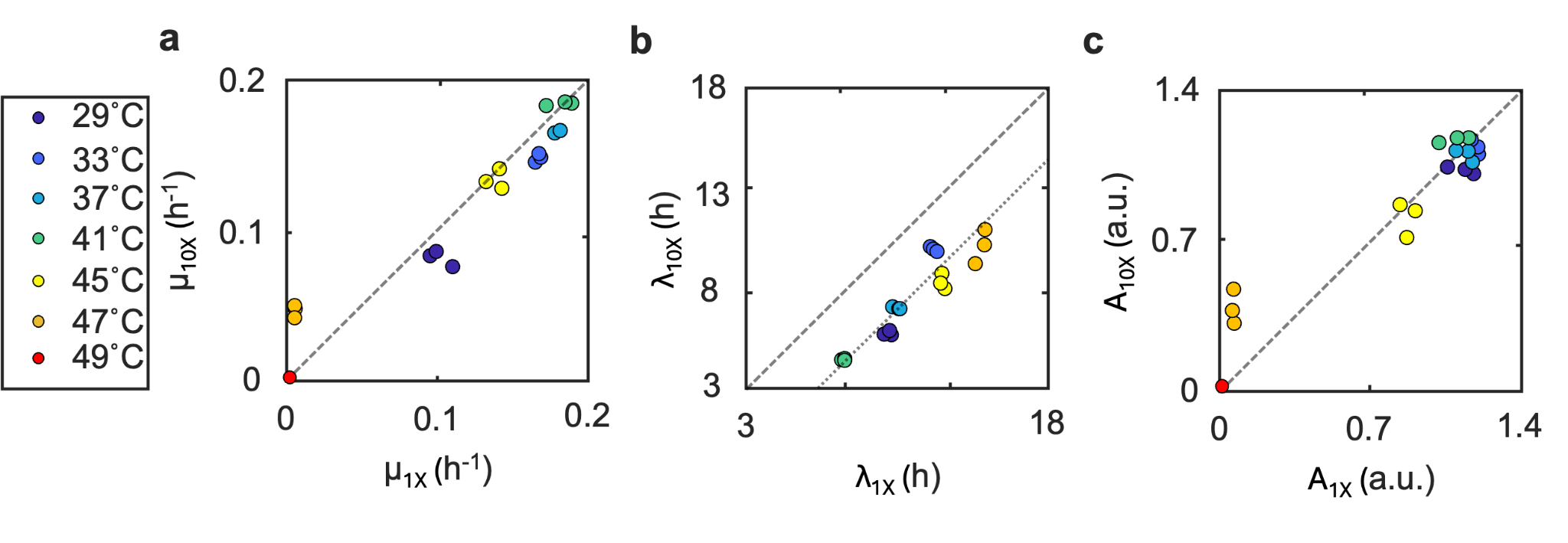

**Fig. S2** Effect of inoculum on thermal response of *L. rhamnosus*. Maximum specific growth rate (**a**), lag phase duration, (**b**) and net growth yield (**c**) of cultures with inoculum sizes of ~10^7^ CFU/mL (represented as 10X) and ~10^6^ CFU/mL (represented as 1X) at different incubation temperatures. Dashed lines indicate points where values are similar between inoculum sizes; in (**b**), the finer dashed line highlights the apparent homogeneous shift of the lag-phase data.

#### E. Viability of thermally stressed cultures

The OD600 remained constant during the 24 h incubation period at 49 °C (and 47 °C with limited change in OD600 value in 1X inoculum). No growth was observed in the cultures at 49 °C. We assessed the viability of cells after 18 h incubation using phase-contrast images of 10X inoculum which revealed a limited number of viable cells in the culture (**Fig. S7(a)**).

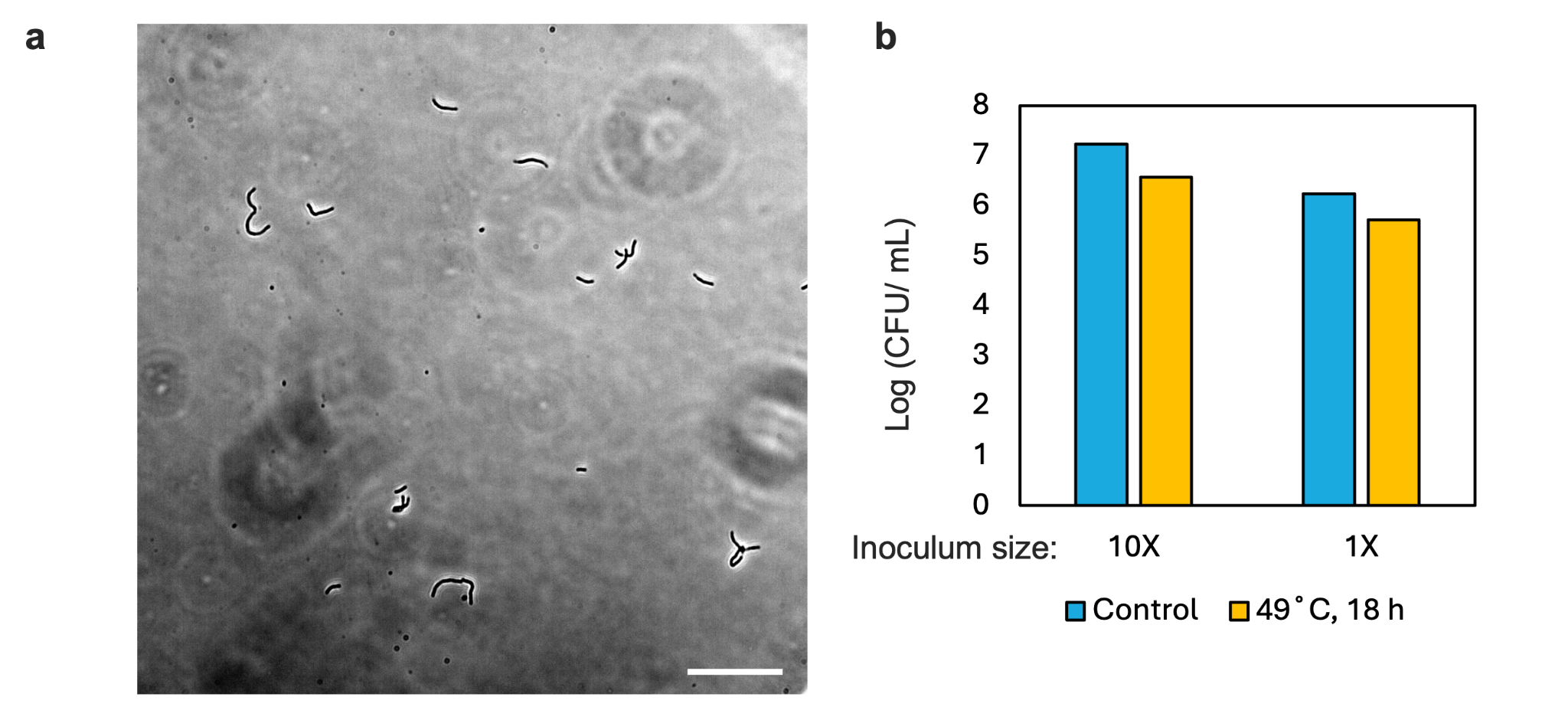

**Fig. S3** Cell viability after incubation at 49 °C for 18 h. (a) Phase-contrast microscopic image of the cells; scale bar is 50 µm. (b) Colony-forming units per mL (CFU/mL) of cultures. Control is the culture before incubation.

To assess the viability of cells, the cells were plated on MRS-agar plates and incubated at 37 °C for 48 h.To minimize the evaporation of medium, we sealed the 96-well plate with parafilm. Note that we considered the evaporated volume of the medium in the calculation of final cell density. We compared the cell densities (CFU/mL) between the initial culture and the corresponding heat-treated sample (**Fig. S7b**). Cells retained their reproductive capacity after 18 hours of incubation at 49 °C; however, the cell density was reduced by 0.7 log. We repeated the experiment with a 10-fold diluted inoculum and obtained a cell density reduction of 0.5 log. Therefore, *L. rhamnosus* cells were able to survive the 49 °C heat treatment, though with a ~1-log reduction in recoverable cell density; and the survivability does not have strong dependency on the initial inoculum size.

### Note 2. Design and fabrication of microfluidic system

#### A. Microfluidic channel network design

The cell and metabolites transport between chambers was controlled through a continuous serial dilution unit where 90% (volume flow rate) of the outgoing stream which contains cells and metabolites is purged and the remnant 10% is diluted with fresh incoming medium before entering the next chamber. Note that we kept the incoming flow rate to each chamber constant to maintain similar hydrodynamic conditions for all chambers (similar transport properties). The minimum mixer channel length to achieve complete mixing for small molecules (Diffusion coefficient of $D={10}^{-10}\frac{m^{2}}{s}$) in a channel of width W = 200 µm and height H = 50 µm under a flow rate of Q_0_ = 160 µL h^–1^ is calculated by

$l_{mix}=\frac{H*Q_{0}}{W*D}$	(4)

Implementing values, we obtain the minimum mixing length of 34 mm (= L_8_ = L_10_ = L_14_ in **Fig. S4**).

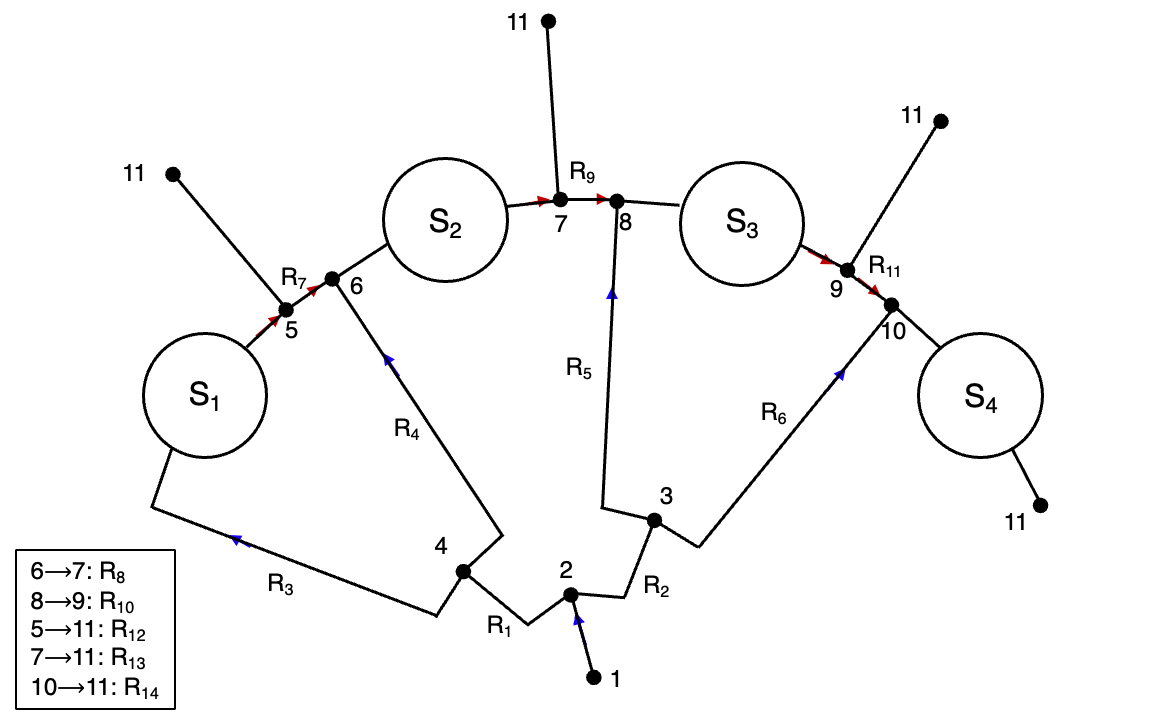

**Fig. S4** Microfluidic network design. Each dot shows a node connected by arrows showing the direction of fluid flow.

The flow distribution in the streams is controlled by changing the hydraulic resistance of large aspect ratio (W/H >>1) rectangular channels (Eq. 11).

$R=\frac{12\mu L}{wh^{3}}$	(5)

Keeping channel height and width constant (H = 50 µm, W = 200 µm), we adjusted channels’ length to achieve the required flow rates in each stream. The following shows the channel lengths that are known from the initial design are L_1_ = L_2_ = 8136 mm, L_4_ = 21596 mm, L_7_ =L_9_ = L_11_ = 6692 mm, L_8_ = L_10_ = 36859 mm. Then, we calculate the rest of the channel length as follows: L_14_ = 41698 mm, L_13_ = 83396 mm, L_12_ = 125094 mm, L_3_ = 20106 mm, L_5_ = 64198 mm, L_6_ =105896 mm.

#### B. Microfabrication

We fabricated the chip by using the conventional soft lithography technique with modifications (Xia and Whitesides, 1998). The high ratio between the chamber height (4 mm) and the channel height (50 µm) makes it impossible to use traditional photo-lithography techniques to manufacture the negative mold of microfluidic devices. The negative mold of channels were created on a silicon wafer by patterning SU-8 (KAYAKU Advanced Materials) using a masked UV lithography technique. Cylindrical pieces with dimensions of chambers were fabricated using Computer numerical control (CNC) milling from polymethyl methacrylate (PMMA) and glued to another silicon wafer. Two molds were then casted with polydimethylsiloxane (PDMS). The two PDMS pieces were then aligned and plasma bonded. The thickness of the PDMS on the chamber floor was kept constant at 2 mm to minimize the heat resistance between the heat source and chamber contents. The final channels design and chambers configuration is discussed and shown in the next section.

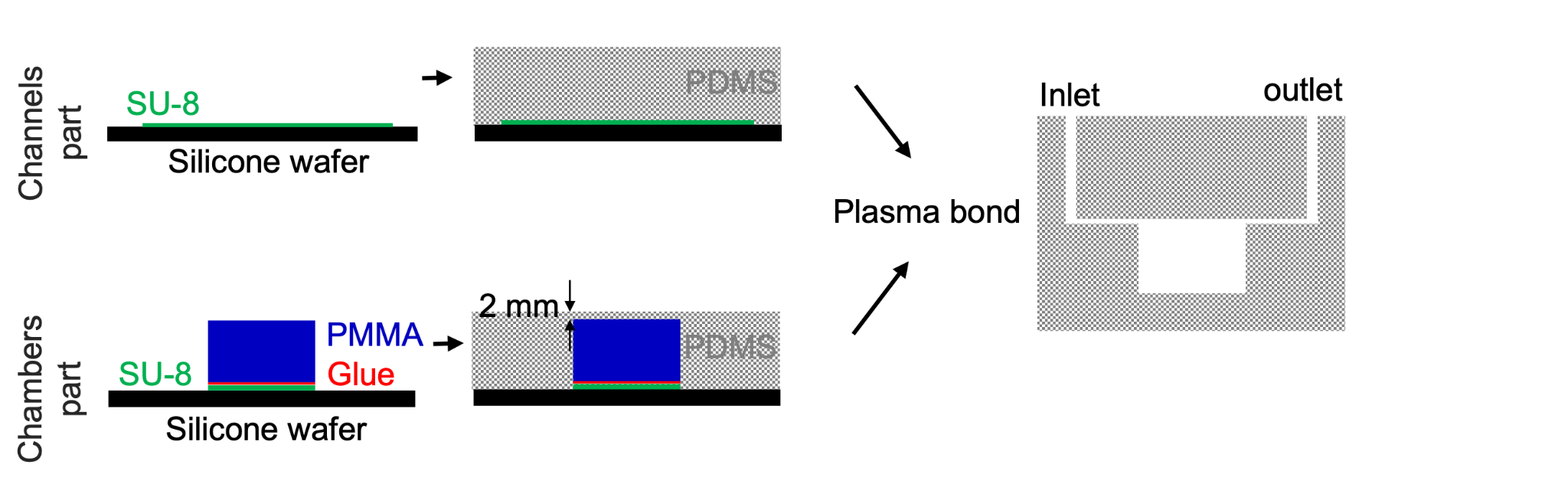

**Fig. S5** Fabrication of the multiscale microfluidic device. The compartments of chambers and channels are manufactured separately from PDMS and plasma-bonded to form the final micro-bioreactor.

#### C. Electrical circuit design for microheaters

To regulate temperature across the four microchambers, Peltier thermoelectric modules ( Same Sky, OR, USA) were used to establish localized heating zones on the µ-Therm platform (**Figure S6a**). These thermoelectric modules use the Peltier effect, converting electrical current into a temperature gradient between their surfaces (Drebushchak, 2008). To ensure efficient heat dissipation, the cold side of each module was coupled with an aluminum heat sink and actively cooled using a 12V DC brushless fan (Sunon Fans). Temperature measurements were facilitated by a thermistor integrated within the thermoelectric module. To optimize thermal conductivity and ensure precise sensing, the thermistor was secured using a thermally conductive adhesive (MG Chemicals, Burlington, ON, CA). This arrangement enabled accurate temperature monitoring and enhanced heat transfer efficiency, maintaining consistent thermal conditions in the microchambers.

**Fig. S6** Electrical design of µ-Therm system. (a) Schematic illustration of the localized thermal heating system, highlighting its key structural components: (1) a custom-designed 3D-printed fr by amework for system integration, (2) thermoelectric Peltier module, (3) thermistor, (4) conductive filling for efficient heat transfer, (5) heat sink, and (6) cooling fan. (b) Electrical circuit diagram of the µ-Therm system, comprising thermoelectric heating modules (gray pads), thermistors (RT1–RT4) for temperature sensing, controllers for real-time regulation, and switches (SW1–SW3) for operational control. The relay controllers are powered by a 12 V supply, with the input voltage to the Peltier modules precisely regulated at 1.4 V to achieve stable and efficient thermal performance.

The electrical circuit comprises four Peltier modules, each regulated by proportional controllers that modulate the electrical current supplied to the thermoelectric modules at 1.4 V via a 12 V DC relay switch (**Figure S6b**). The input voltage to the Peltier modules is provided by a centralized power source, selected in accordance with manufacturer specifications to ensure optimal performance. To streamline operation, all components are connected in parallel to the primary power source, ensuring uniform current distribution across the system. Individual switches are incorporated into the circuit design, enabling independent control over each component for precise system management and operational flexibility.

### Note 3. Thermal and hydrodynamic characterization

#### A. Mathematical modeling of heat transfer to microchambers

To investigate heat transfer from the heater to the microchamber, a simplified model of heat conduction in solids and liquids was used (**Figure S3**).

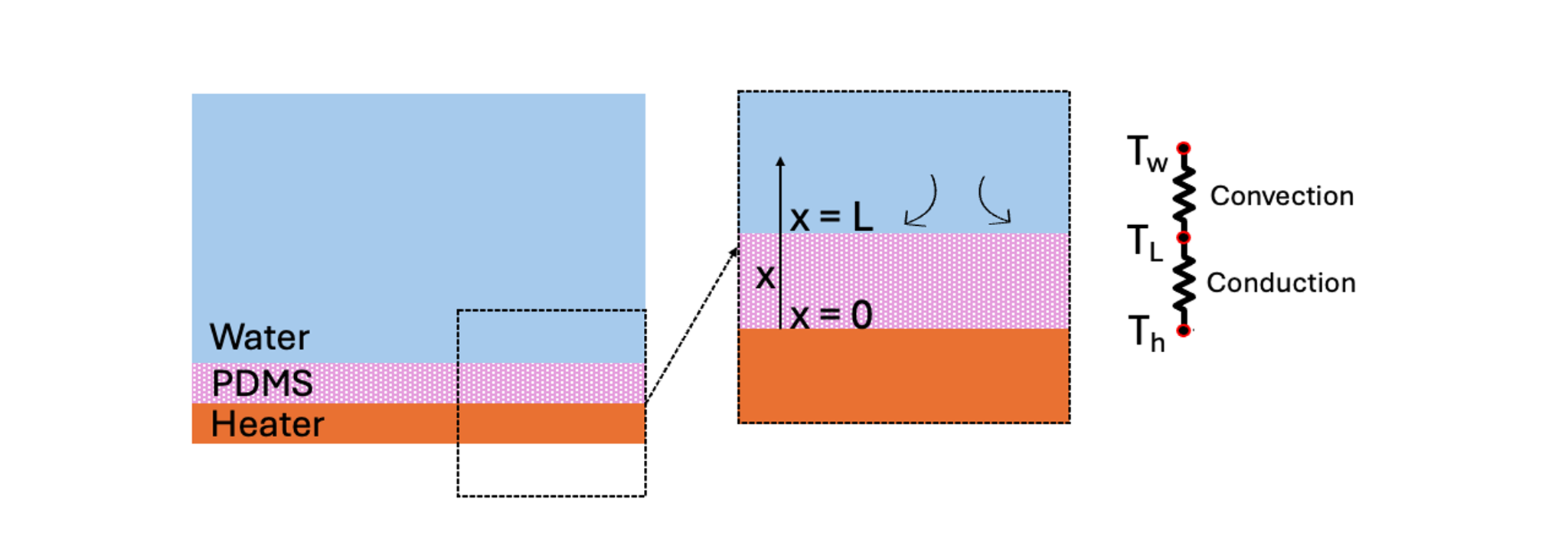

**Figure S7.** Schematic representation of the heat transfer model for the microchamber system. Heat propagates from the heated surface of the µ-Therm device to the thin PDMS layer of the microchamber via conduction. Subsequently, it transfers to the growth medium, modeled as water, primarily through convection. This dual-mode heat transfer mechanism ensures precise thermal control, critical for biological processes within the microchamber.

The system comprises a polydimethylsiloxane (PDMS) layer of thickness L, where heat transfer is governed by the diffusion equation (1):

$\rho c\frac{\partial T}{\partial t}=\nabla.(k\nabla T)$	(6)

where ρ, c, and k are density, specific heat, and thermal conductivity of PDMS. By assuming one-dimensional heat conduction in x direction in PDMS, **Equation 1** is simplified to **Equation 2** describing spatiotemporal temperature change in PDMS.

$\rho c\frac{\partial T}{\partial t}=k\frac{\partial^{2}T}{\partial x^{2}}$	(7)

At PDMS-heater interface x = 0, the temperature T_s_(x,t) varies periodically– due to proportional closed-loop controller– and, by applying Fourier series expansion, its oscillation can be considered as T_s_(0,t) = T_h,0_ + A cos(ωt), where T_h,0_ is the average temperature of the heater surface, A is the amplitude of temperature oscillation, and ω is the angular frequency of oscillation. At water-PDMS interface x = L_1_, there is a conductive-convective heat transfer condition ensuring that there is continuity of temperature and heat flux at this interface, considering both heat conduction in PDMS and convective heat transfer between PDMS and water. This boundary condition can be assumed as $-k\frac{\partial T}{\partial x}|_{x=L}=h(T\left( L,t \right)-T_{w})$, where h is the convective heat transfer coefficient between PDMS, and water and T_w_ is the water temperature. Using steady periodic solution approach and considering the boundary conditions, T(L ,t) can be expressed as:

$T\left( L,t \right)=T_{h,0}+A_{L}cos(\omega_{L}t-\phi)$	(8)

where A_L_, ω_L_, and ϕ are amplitude of temperature oscillation at x = L, frequency of oscillation at x = L, and phase shift determined by phase lag due to distance L and convective heat transfer. Frequency of temperature oscillation at x = L is same as the heater temperature oscillation; however, it has a phase shift of $\phi=(\frac{hL}{k})$. The amplitude of oscillation decreased by a factor of $\left( 1+\left( \frac{hL}{k} \right)^{2} \right)^{-0.5}$. For h calculation, we assumed a natural convection on the chamber floor since the liquid flow is laminar on the surface h = k/L_c_×Nu. Nu is Nusselt number can be correlated with Grashof number (Gr) and Prandtl number (Pr) for a flat horizontal surface by(Welty et al., 2014)

$Nu=0.59 \left( Gr\times Pr \right)^{0.25}$	(9)

$Gr=\frac{g\beta\left( T\left( L,t \right)-T_{w} \right)}{\nu^{2}}$	(10)

$Pr=\frac{\nu}{\alpha}$	(11)

L_c_ is characteristic length calculated by L_c_ = a/p, where a is area of heating and p is perimeter of PDMS surface. L_c_ is 0.0019 m for the microchamber. By substituting parameter values from **Table S1** to **Equations 2 through 6** and assuming T(L,t) = 50 ˚C, we get h = k_f_/L_c_×Nu = 1396 W.m^–2^.K^–1^.

**Table S2.** Parameters and values considered for mathematical modeling of heat transfer to microchamber

| **Parameter** | **Description** | **Value** |
| --- | --- | --- |
| L | Thickness of PDMS in microchamber floor (mm) | 2 |
| ρ | Density of PDMS (kg.m^-3^) | 970 |
| c | Specific heat of PDMS (J.kg^-1^.K^-1^) | 1200 |
| k | Thermal conductivity of PDMS (W.m^-1^.K^-1^) | 0.2 |
| T_h,0_ | Average temperature of heater surface (˚C) | 50 |
| A | Amplitude of temperature oscillation on heater surface (˚C) | 5 |
| ω | Angular frequency of temperature oscillation on heater surface (Hz) | 0.06 |
| T_w_ | Water temperature (assumed to be constant) (˚C) | 22 |
| g | Acceleration due to gravity (m.s^-2^) | 9.8 |
| β | Thermal expansion coefficient of water at 22˚C (K^-1^) | 3.84×10^-4^ |
| c_f_ | Specific heat capacity of water (J.kg^-1^.K^-1^) | 4184 |
| k_f_ | Thermal conductivity of water at 22˚C (W.m^-1^.K^-1^) | 0.628 |
| ν | Kinematic viscosity of water at 22˚C (m^2^.s^-1^) | 0.66×10^-6^ |
| α | Thermal diffusivity of water at 22˚C (m^2^.s^-1^) | 0.15×10^-6^ |
| a | Effective surface area through which heat transfer occurs (m^2^) | 4.5×10^-5^ |
| p | Perimeter of PDMS surface being heated (m) | 23.910^-3^ |
| ρ_f_ | Density of water (kg.m^-3^) | 1000 |
| V_f_ | Volume of water in microchamber (m^3^) | 180×10^-9^ |

By substituting the value of parameters listed in **Table S2** to **Equation 3**, we can estimate A_L_ as 0.5 ˚C. This gives an amplitude ratio (AR = A_L_/A) of 0.1. To assess this experimentally, we measured the temperature of a single microchamber of the TALE chip containing MRS broth using a thermocouple placed on the chamber floor over time and recorded the temperature on the controller screen. **Figure S8** compares the temperature, normalized by maximum temperature in the period of oscillation, shown on the controller and temperature measured inside the chamber. The calculated AR from experiments is 0.01 which is within one order of magnitude smaller than theoretically calculated value. Hence, oscillation of heater temperature oscillation will not have a big impact on the temperature inside the microchamber where the bacteria will be grown.

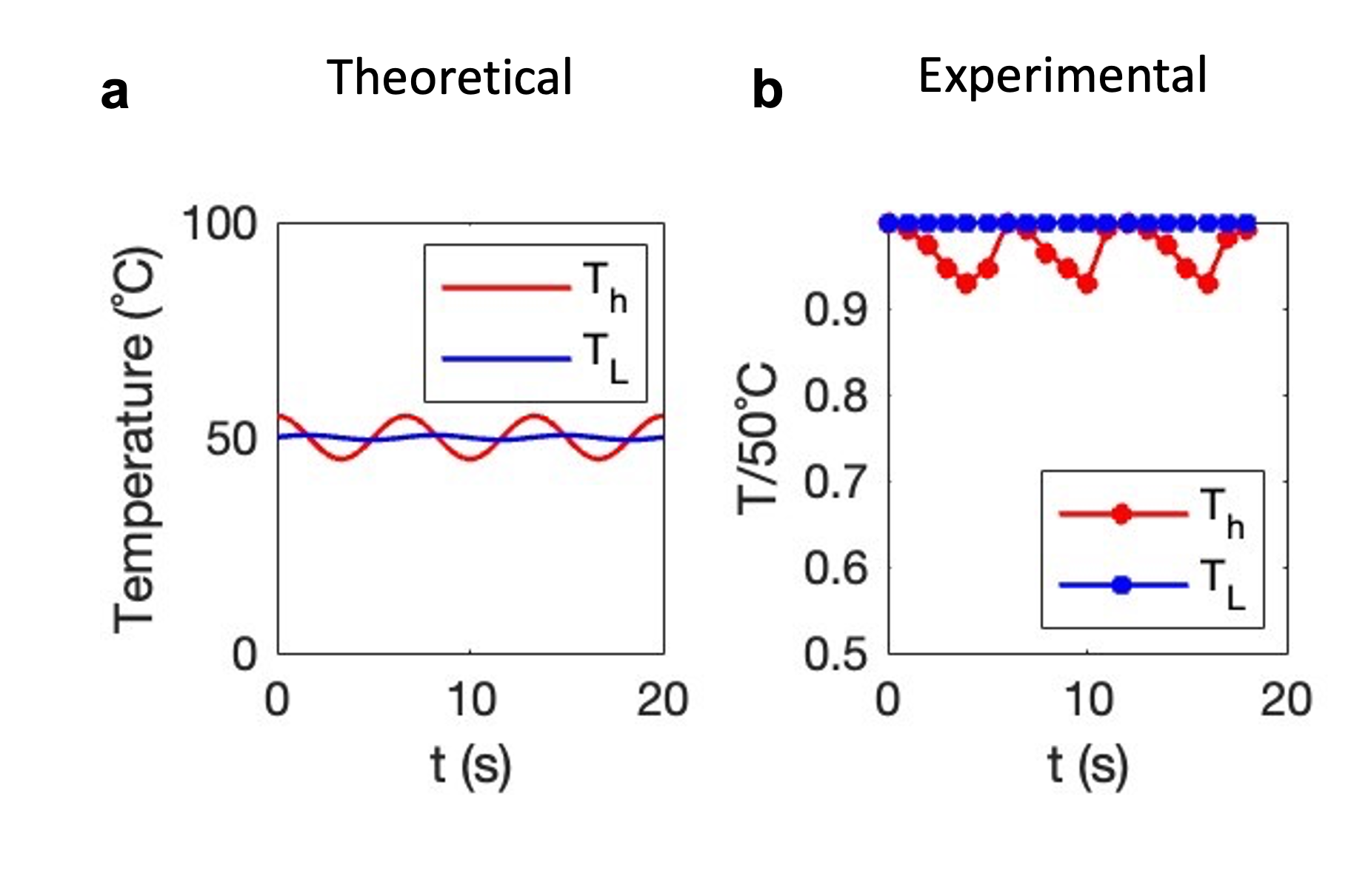

**Fig. S8.** Comparing oscillation of temperature on the µ-Therm heater T_h_ and inside the microchamber T_L_ resulted from (a) theoretical modeling and (b) experimental investigation.

#### B. Residence time of cells in chambers

We aim to simulate the transport of incoming bacterial cells to a cylindrical chamber. The micro-chamber is a cylinder with a diameter of 7.6 mm and height of 4 mm resulting in a total volume of 180 µL. Refreshing medium flow is supplied through microchannels of height 50 µm and width of 200 µm (**Fig. S1a**). Bacteria are modeled as spherical particles with a diameter of 1 µm (analogous to the bacteria size) being transported within a chamber. We used COMSOL Multiphysics (version 6.0) to simulate the transport of bacteria to the chamber. In the inlet, 1000 particles were released from the inlet. The flowrate of the inlet stream was in the range of 10 µl.hr^–1^ and 300 µl hr^–1^. Outlet boundary condition was set on zero static pressure. The flow field was calculated for by a stationary study. The fluid is considered to be incompressible with a density of 1000 kg.m^–3^ and viscosity of 1×10^–3^ Pa.s. A Lagrangian particle tracking model was used to predict the particle trajectory over time. The particles were subjected to viscous drag force and gravity force to consider both the hydrodynamic transport and sedimentation of particles. The large ratio of chamber to channel height (= 4 mm /0.05 mm = 80) creates a strong expansion-contraction zone for particles which restricts the transport (**Fig. S1b**). The residence time (𝜏) was calculated from particle location data obtained from the simulation and analyzed using a Python script. Increasing the flow rate reduced the residence time, which plateaued at approximately 20 min. The data is well-fitted to a logistic function (**Eq. 1**)

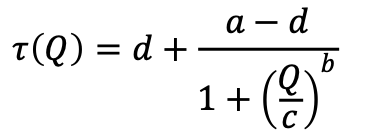
(12)

where a = 20, b = -1.4, c = 7.9, and d = 1.0 are constants.

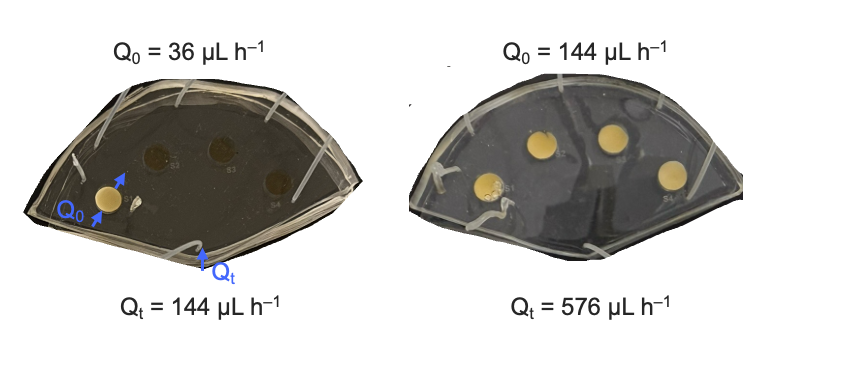

**Fig. S9**. Effect of refreshing medium flow rate on bacterial growth in the microfluidic device. Photos indicate the growth of *L.rhamnosus* in the microfluidic device after 48 h incubation of all chambers at 37 ºC. Minimum flow rate (36 µL h^–1^) is insufficient for continuous cultivation.

### Note 4. Biomass formation and dynamics in microchambers

Tracking the pixel intensity of individual microchambers provides a detailed temporal profile of growth initiation and biomass formation within the confined environment of a microfluidic device. For instance, microchamber S2 exemplifies distinct phases of bacterial proliferation, characterized by an observable lag phase followed by exponential growth. Notably, growth initiation is detected approximately 12 h after inoculation, emphasizing the dynamic adaptation and development of the bacterial population within the microchamber (**Figure S10**).

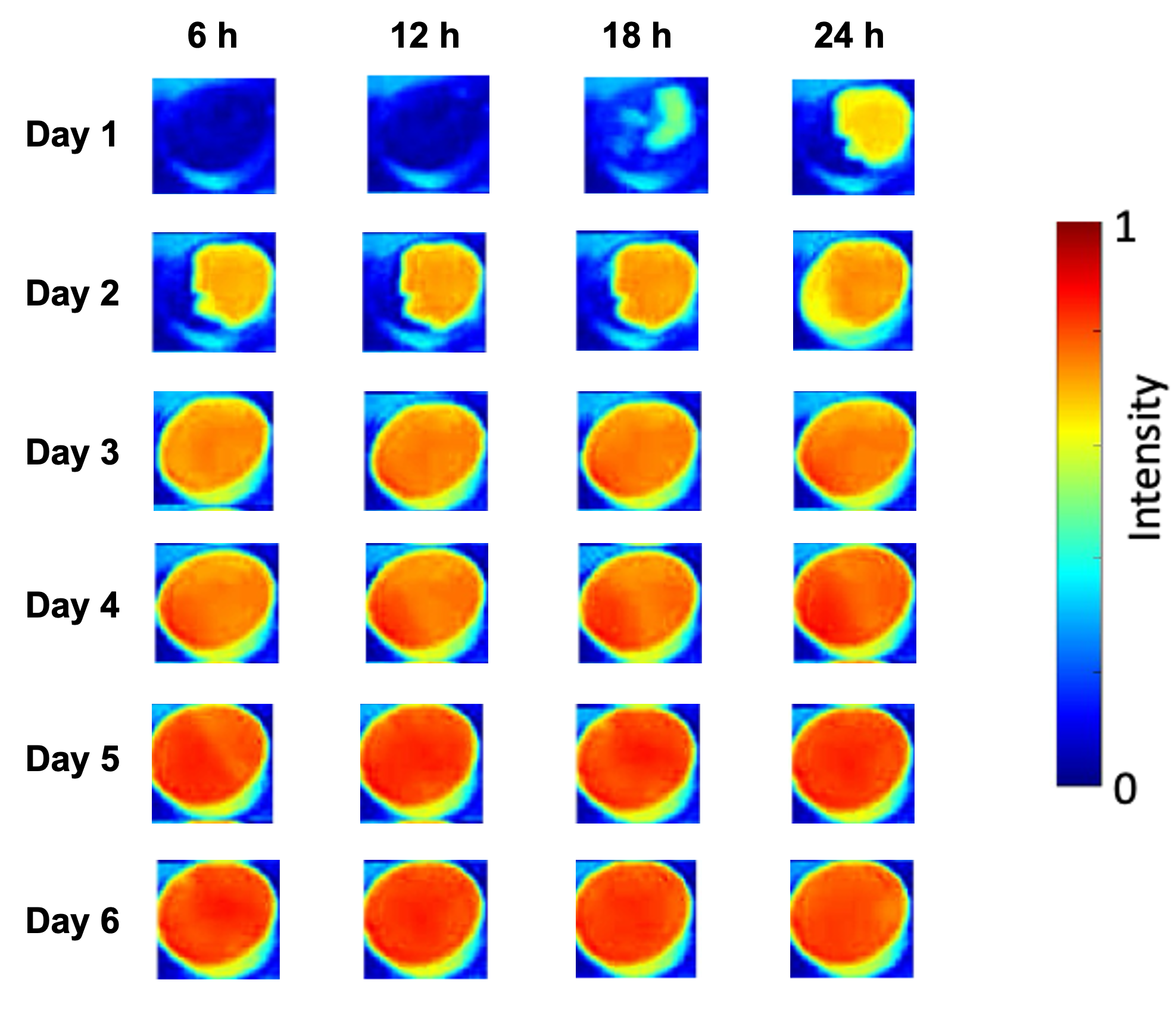

**Fig. S10.** Intensity profile of microchamber S2 over the time course of 6 d (144 h). Higher intensity indicates the amount of biomass generated in the microchamber.

We identified four distinct phases of bacterial growth within the microchamber: (i) the planktonic phase, marked by the presence of free-floating cells; (ii) the stationary biomass phase, characterized by localized cell aggregation and growth; (iii) horizontal growth, defined by lateral proliferation along the microchamber surface; and (iv) direct transport, driven by mechanical pressure resulting from upstream biomass accumulation. These phases are schematically illustrated in **Figure S11**, highlighting the role of natural convection in shaping spatial growth dynamics.

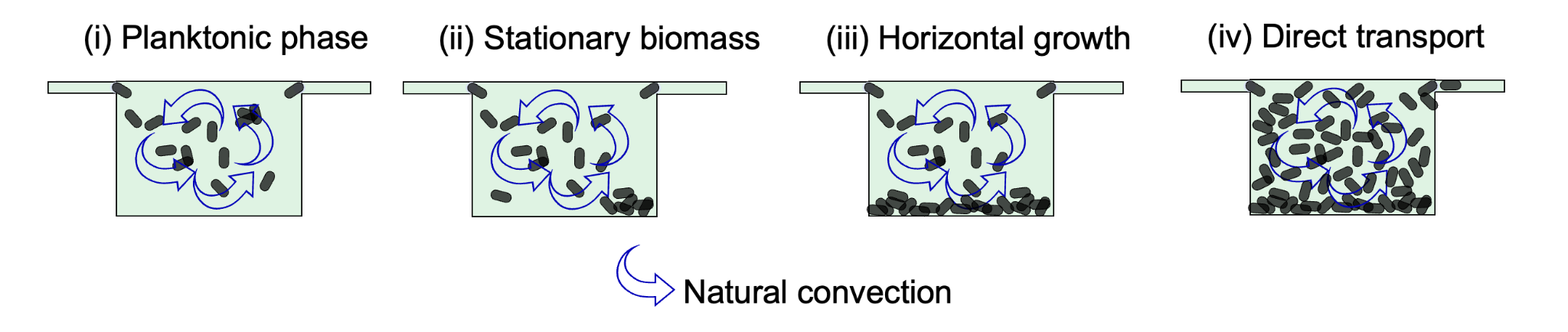

**Fig. S11**. Phases of bacterial growth in the microchamber: (i) Planktonic phase with free-floating cells, (ii) Stationary biomass phase with settled aggregates, (iii) Horizontal growth with lateral expansion, and (iv) Direct transport driven by upstream biomass pressure. Blue arrows depict natural convection patterns.

Heating from beneath the microchamber induces natural convection, driving fluid circulation within the microchamber. Using COMSOL Multiphysics software (version 4.5), the velocity field at the microchamber’s mid-cross-section by modeling laminar flow and heat transfer under a flow rate of 150 µl h^–1^ and a floor temperature of 50 °C was simulated. The interplay of external fluid flow and natural convection results in fluid circulation (**Fig. S12a**) and a homogeneous temperature profile (**Fig. S12b**). Due to the microchamber walls, the lowest fluid velocities, below 1.5 µm s^–1^, are observed near the edges. These velocities are insufficient to transport spherical particles of 1 µm diameter, allowing biomass to settle in these low-velocity zones. These regions correspond to the initiation of stationary phase biomass growth, as indicated by the chamber intensity profile (**Fig. S12c**).

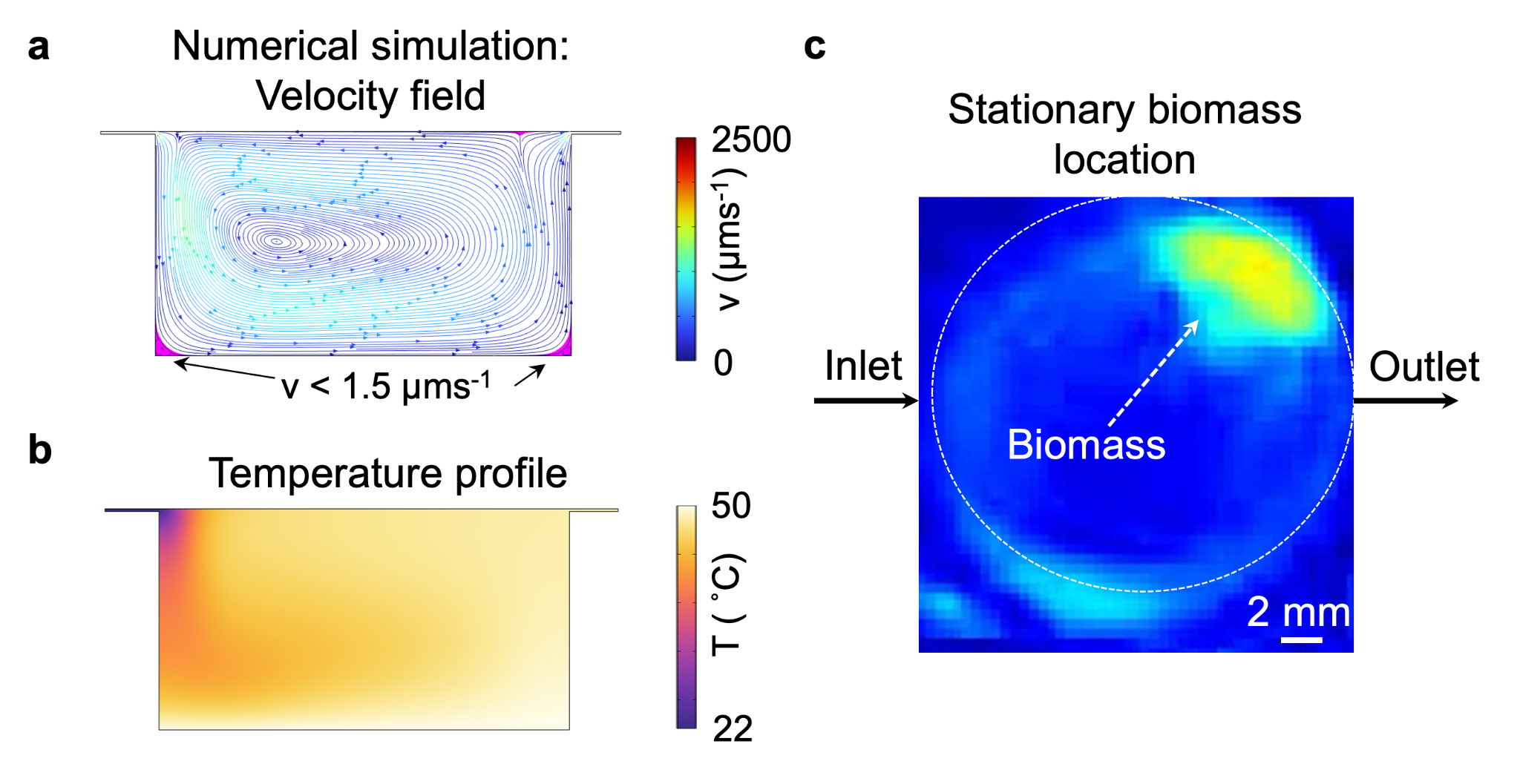

**Fig. S12**. Numerical simulation and experimental intensity profile of biomass settlement in a microchamber. (a) Velocity field and (b) temperature distribution in the mid-cross section of a microchamber with dimensions consistent with those used in this study. The simulations, conducted in COMSOL Multiphysics, assume water as the working fluid, with a floor temperature of 50 °C and an inlet fluid flow rate of 150 µl h^–1^. Cyan-colored regions indicate areas of potential biomass settlement, where stationary biomass growth is expected to initiate. (c) Intensity profile of microchamber S4 during the stationary biomass phase, demonstrating a biomass settlement pattern that aligns with the numerical predictions.

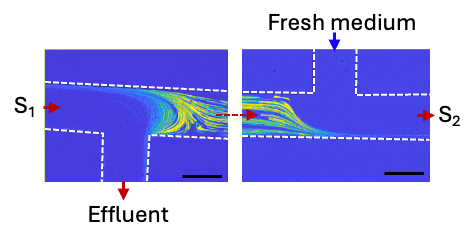

**Figure S13.** Continuous serial dilution (CSD) unit operation between chambers S_1_ and S_2_. Overlay of 20 frames of cell motion at Q = 144 µL h^–1^ captured using a phase-contrast microscope. The image is colorized according to pixel intensity. Brighter colors show the cell track.

**Table S3**. Optical density (OD600) of cultures extracted from the experiment on the thermal stress landscape.

| **Sample** | **100×OD600** |
| --- | --- |
| S1 | 0.219 |
| S2 | 0.150 |
| S3 | 0.124 |
| S4 | 0.124 |

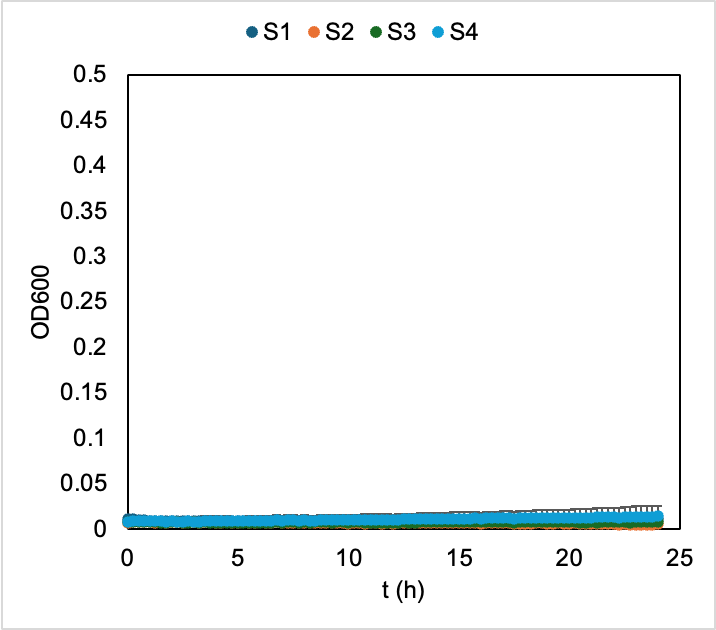

**Fig. S14**. Growth curves at 49 ºC of the on-chip grown cultures in the periodic thermal landscape.

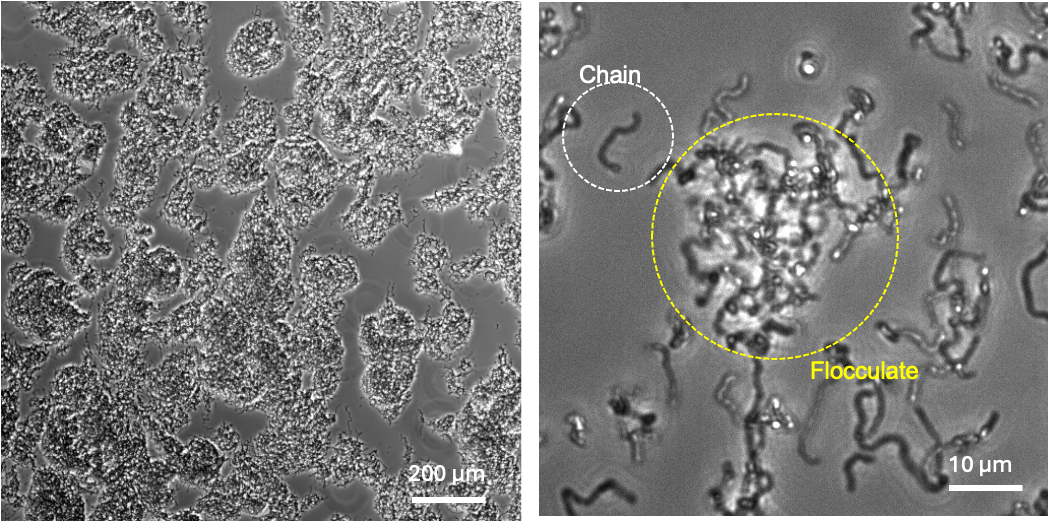

**Fig. S15.** Phase-contrast microscopy images of cells collected from the S1 chamber after 48 h of incubation. The cells exhibit pronounced chain formation, with these chains further aggregating into flocculated clusters.

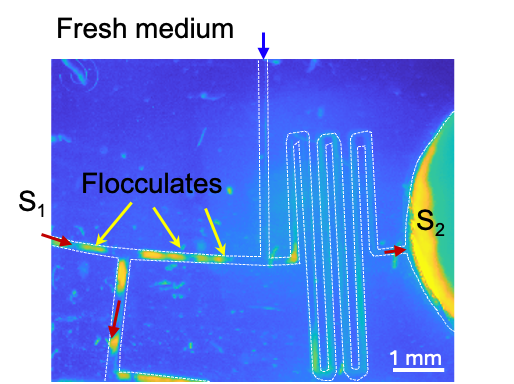

**Fig. S16**. False-colored image of the CSD unit showing the transport of fluctuates between microchambers. Yellow color indicates flocs.
